# Cross-talk between engineered *Clostridium acetobutylicum* and *Clostridium ljungdahlii* in syntrophic cocultures enhances isopropanol and butanol production

**DOI:** 10.1101/2025.07.23.666411

**Authors:** Jonathan K. Otten, John D. Hill, Noah B. Willis, Joseph Dougherty, Andrew Dalton, Eleftherios T. Papoutsakis

**Affiliations:** Department of Chemical and Biomolecular Engineering, University of Delaware, Newark, DE, USA; Delaware Biotechnology Institute, University of Delaware, Newark, DE, USA; Department of Biological Sciences, University of Delaware, Newark, DE, USA

**Keywords:** Clostridium acetobutylicum, Clostridium ljungdahlii, coculture, isopropanol, butanol, consortia, metabolic flux analysis

## Abstract

There is a need for efficient and sustainable production of essential chemicals such as isopropanol and butanol – the focus of this study – from renewable sugar feedstocks. Microbial fermentations use glycolysis, and as result, a third of the sugar carbon is lost to CO_2_ through pyruvate decarboxylation to acetyl-CoA, the starting intermediate for the biosynthesis of most microbial metabolites. In nature, microbes exist in syntrophic consortia, allowing for mutually-beneficial interactions, the production of novel products and the realization of novel benefits – including better carbon conservation – not seen in monocultures. Here, for increased acetone production, we transformed *Clostridium acetobutylicum* with a plasmid (p95ace02a) expressing a synthetic acetone pathway made up of four native genes. This engineered *C. acetobutylicum* was cocultured with *Clostridium ljungdahlii* to capture the waste CO_2_ and H_2_ generated due to glucose catabolism by *C. acetobutylicum*, and to convert acetone into isopropanol. We examined the impact of starting cell densities, the gas atmosphere (N_2_, H_2_, or H_2_/CO_2_) and coculture species ratios (using a recently-developed RNA-FISH flow cytometric assay) on metabolite production, yields and sugar-carbon utilization. Metabolic flux analysis identified the complex patterns by which the two species alter each other’s metabolism in a cell-density and gas-atmosphere dependent manner. For example, *C. ljungdahlii* activated the dormant acetate uptake in *C. acetobutylicum*, while coculture density dramatically impacted species ratios, electron management, and *C. ljungdahlii’*s H_2_ utilization. We achieved exceptionally-high concentrations of our desired products – 246 mM isopropanol and 148 mM butanol – in 64 hours, with about 85% of the production occurring before 32 hours. We reached maximum productivities of 13.9 mM isopropanol/hour and 10.4 mM butanol/hour with 0.9 mol alcohol produced per mol of sugar consumed. Total product yields reached 84.7% on a C-mol basis, versus 65.6% that can be reached in a *C. acetobutylicum* monoculture.

## Introduction

As the world moves towards more sustainable means of chemical manufacture, non-petroleum-based pathways for the production of essential chemicals such as isopropanol, butanol, ethanol, and acetone (IBEA) are needed. In the past, microbial fermentation with *Clostridium acetobutylicum* (*Cac*) has been used to produce butanol, ethanol, and acetone (BEA) (Gabriel and Crawford 1930, Gibbs 1983, Lutke-Eversloh and Bahl 2011, Adams 2017). However, these processes were rendered economically uncompetitive after World War II due to the rise of petroleum-based processes and to their own inherent inefficiency: in *Cac* monoculture fermentations, a third of the sugar carbon is lost as CO_2_ (Charubin and Papoutsakis 2019).

These marketplace inefficiencies do not blunt some of the main advantages of microbial fermentations with *Clostridia*, and new biotechnological advances can engineer greater control over the microbial organisms’ genetics, the operation and analysis of microbial fermentation, and the pairing of organisms to capture waste CO_2_ and to expand the metabolic space (Lutke-Eversloh and Bahl 2011, Charubin, Bennett et al. 2018, Charubin and Papoutsakis 2019, Jiang, Wu et al. 2023). *Cac* is an appealing model industrial microorganism because it has a broad and powerful substrate utilization system and primary metabolism and because it exhibits high electron and carbon fluxes and distinctive systems for generating and shuffling electrons (Charubin, Bennett et al. 2018). Metabolic engineering has enabled the design of plasmids that produce butanol, ethanol, and acetone at greater concentrations and purities (Lee, Jang et al. 2009, Sillers, Al-Hinai et al. 2009, Lutke-Eversloh and Bahl 2011), added isopropanol production capabilities (albeit at low concentration) (Lee, Jang et al. 2012, Dusseaux, Croux et al. 2013), and used antisense RNA strategies to change the balance of metabolite production (Desai and Papoutsakis 1999).

Here, we augment a *Cac* fermentation with in-situ carbon-capture via coculture with *Clostridium ljungdahlii* (*Clj*), an acetogen that fixes CO_2_ using H_2_ as an electron source. *Clj*, an acetogen, fixes CO_2_ using H_2_ as an electron source, making it an ideal coculture partner for *Cac*, which generates both CO_2_ and H_2_ as byproducts of glucose fermentation (Jones, Fast et al. 2016, Charubin and Papoutsakis 2019). Our lab has already demonstrated that, in *Cac*/*Clj* cocultures, *Clj* expands the metabolic space by enabling the production of isopropanol and 2,3-butanediol, which neither *Cac* nor *<u>Clj</u>*is capable of on its own. *Clj* also produces acetate, which can be reassimilated by *Cac*. Previous research has demonstrated that direct physical contact aids this metabolic coupling (Charubin and Papoutsakis 2019); TEM tomography shows a distinct physical contact phenotype by which *Clj* makes contact with *Cac* at its poles, forming fusion pairs (Charubin, Modla et al. 2020). We have also confirmed that cytoplasmic proteins, RNA, and DNA are transferred between cells (Charubin, Hill et al. 2024). This study builds on that foundation by incorporating an engineered *Cac* strain alongside *Clj* in pH-controlled bioreactors to maximize IBEA productivity.

The studies of cocultures across *Clostridial* species are increasingly relevant. Cocultures of *Cac* and *Clostridium kluyveri* (Otten, Zou et al. 2022), or *Clj* and *C. kluyveri* (Richter, Molitor et al. 2016), have been used to produce 6-8 carbon compounds such as hexanoate. *Clostridium beijerinckii* has also been partnered with *Propionibacterium freudenreichii* (Hocq and Sauer 2022) and *Bacillus subtilis* (Cui, He et al. 2020) to produce BuOH, IPA, and acetone. We aim to produce IBEA more quickly and at higher concentrations while utilizing coculture flux analysis (Papoutsakis 1984, Desai, Nielsen et al. 1999) and RNA-FISH (Hill and Papoutsakis 2024) to analyze how coculture interactions change the metabolic fluxes and population ratios of the member organisms.

## Materials and Methods

**Abbreviations:** *Cac: C. acetobutylicum*; *Clj: C. ljungdahlii;* IPA: isopropanol; BuOH: butanol; EtOH: ethanol

### Strain and plasmid construction

The p95ace02a plasmid was described in our prior publication (Seo, Capece et al. 2024). Briefly, it contains the *Cac ctfA* and *ctfB* genes under the control of the *pta* promoter from *Clj*, and the *Cac thl* and *adc* genes under the control of the Pthl^sup^ promoter (Streett, Kalis et al. 2019), the ColE1 origin of replication, *repL*, Amp^R^, and MLS^R^. It was constructed using the NEBuilder HiFi DNA Assembly kit (New England Biolabs) with necessary primers and gBlocks from IDT. The PCR fragments were amplified with Phusion DNA polymerase (Thermo Fisher Scientific). The plasmid construction was successfully confirmed with colony PCR with Phire DNA polymerase (Thermo Fisher Scientific) and Sanger sequencing.

### Transformation of *C. acetobutylicum*

*Cac* was electroporated with the plasmid p95ace02a as previously established (Mermelstein and Papoutsakis 1993) and recently detailed (Seo, Capece et al. 2024).

### Media composition

Luria-Bertani (LB) medium was used for the propagation of *E. coli*. For plates, 15 g/L Select Agar (MilliporeSigma, MA, USA) was added. *Cac* cells were grown in T-CGM-G as previously described (Charubin and Papoutsakis 2019, Otten, Zou et al. 2022). 20 mL/L of potassium phosphate buffer consisting of 100 g/L of KH_2_PO_4_ and 125 g/L of K_2_HPO_4_ adjusted to a pH of 6.8 was used instead of 10 mL/L. *Clj* cells were grown in this T-CGM-G medium with the glucose removed, termed T-CGM-N. Bioreactors and serum bottles used T-GCM-G with varying levels of glucose. When needed, additional glucose was supplemented with a 600 g/L stock. 1.8 M NaOH and 1.5 M HCl were used for pH adjustment. To prepare the bioreactors, they were autoclaved with T-CGM-G before the phosphate buffer, glucose, fructose, Wolfe’s Vitamins, antibiotics, and antifoam were added. These additions were all injected into the bioreactors after the reactors cooled. The antifoam was a solution of 10% Antifoam 204 (Teknova). The media was deoxygenated inside the anaerobic chamber for at least 24 hours or rapidly sparged on a degassing rig with N_2_. The bioreactors were also sparged with N_2_ for at least 2 hrs. When necessary, the following concentrations of antibiotics were used: 100 μg/mL ampicillin,100 μg/mL erythromycin, 50 μg/mL clarithromycin.

### Cell preparation for inoculation

To prepare *Cac* cells for coculture inoculation, single colonies were picked from an agar plate in the anaerobic chamber and transferred to 15 mL conical tubes containing 10 mL of T-CGM-G. The tubes were sealed and heat-shocked at 80°C. The tubes were then transferred back to the anaerobic incubator. Once cooled to 40-50°C, clarithromycin was added and the screwcaps were left loose to allow for gas exchange. Once the cells grew to an OD_600_ of 3-5, they were passaged at 10x, 100x, and 1000x dilution to 50 mL conical tubes or GL-45 glass laboratory media bottles to reach the appropriate volume of cells at an OD_600_ of 3-5. Multiple dilutions were chosen to allow for optimal synchronization of *Cac* cells at mid-exponential phase to *Clj* cells at mid-exponential phase. Once the final inoculum OD_600_ measurements were taken, the cells were concentrated via centrifugation at 8000 x *g* for 8 minutes in 50 mL conical tubes and transferred to 30 mL syringes within the anaerobic chamber for inoculation into the bioreactors.

*Clj* cells were started from glycerol frozen stocks into YTAF-MES media (containing 14 g/L tryptone, 9 g/L yeast extract, 1.4 g/L L-arginine, 10 g/L fructose, and 10 g/L MES) (Cooksley, Zhang et al. 2012). An entire 1 mL tube of this -80°C freezer stock is added to a 160 mL serum bottle containing 25 mL YTAF-MES and clarithromycin. The headspace of the bottle is purged with a gas composed of 80% H_2_ and 20% CO_2_ and pressurized to 22 psi. These bottles are left to grow on a shaker at 37°C. Once the OD_600_ of the *Clj* reaches 0.7-1.0, the cells are passaged to 500-1000 mL serum bottles containing 20% of their total volume of T-CGM-N with clarithromycin. The bottles are likewise purged and pressurized to 22 psi with the H_2_/CO_2_ blend. Once the cells reach an OD_600_ of 0.7-1.0, an appropriate volume of cells are concentrated via centrifugation following the same process as *Cac*. Because *Clj* grows more slowly, the target OD_600_ ratio was 4-8 *Clj* : 1 *Cac*.

### Bioreactor setup and operation

The small-scale bioreactors used in this experiment have been described (Otten, Zou et al. 2022). Briefly, spinner flasks (Chemglass CLS-1400-100) were used with their spinner assembly removed. An active volume of 150-200 mL was used. This covers a pH probe that has been inserted through an open GL-32 cap and secured by a grommet. The other GL-32 port hosted a solvent delivery cap with four ports: sampling, base, sparging, and exhaust. The pH was controlled via networked, remotely-accessible pH controllers (Bluelab). They supplied 1.8 M NaOH to each reactor. Each vessel could be adjusted for sparging or headspace gassing through manipulation of the internal length of the PTFE gassing tube. Individual gas flow rates to each reactor were controlled by flowmeters (Dwyer). The reactors can use any nonflammable gas blend, including pure N_2_, pure CO_2_, or a blend, including 5% H_2_, 10% CO_2_, and 85% N_2_. Gas is exhausted through a tube that terminates in a water-filled flask to allow for visible bubbling and separation for the aerobic atmosphere. The vessels are placed in a large rectangular tub that sits over a bed of stir plates. The stir plates turn magnetic stir bars within each vessel that allow for gentle mixing. The tub is filled with water that is maintained at 37°C by sous vide circulators.

All vessels are first autoclaved with DI water after being bleached and cleaned from prior runs. This water is then discarded, and then fresh T-CGM-G media, sans sugar, phosphate buffer, Wolfe’s vitamins, antibiotics, and antifoam, is added to the bioreactors and autoclaved. These supplements are added once the reactors cool from the autoclave. The pH probes are calibrated and sterilized with a solution of hard water and dilute bleach before being rinsed with DI water and ethanol and inserted into the port grommet. Once fully assembled and checked for leaks, the bioreactors are sparged with N_2_ at 50 cc/min for at least 2 hrs. The vessels are then ready for inoculation with syringes of concentrated cells that have been in the anaerobic chamber.

### Preparation of serum bottles for coculture experiments

To prepare 160 mL serum bottles for coculture experiments, the bottles were autoclaved with foil covering the aperture, passed into the anaerobic chamber, filled with 20 mL of T-CGM-G and concentrated cells, sealed with a sterile rubber stopper and crimped with an aluminum ring. The bottles were then passaged outside of the chamber where they are purged with gas for several minutes and pressurized to 20 psi. Normally, a blend of 80% H_2_ and 20% CO_2_ was used unless the experiment called for manipulation of the headspace composition.

### High-performance liquid chromatography analysis of metabolites

At each timepoint, the pH, OD_600_, and metabolic profile of the fermentations were measured. The metabolic composition of the media was analyzed as described (Carlson and Papoutsakis 2017, Otten, Zou et al. 2022, Seo, Capece et al. 2024). An HPLC system (Agilent) was used with an Aminex HPX-87H column (Bio-Rad) and a running buffer of 5 mM H_2_SO4 at 0.5 mL/min. Additionally, to monitor the approximate glucose concentration of the fermentations, a refractometer was used (Milwaukee Instruments). The refractometer values are not reported in this work as the HPLC measurements are more accurate, but the refractometer results informed the supplemental addition of any requisite glucose before the HPLC samples could be processed.

### RNA-FISH sample prep and flow-cytometric analysis

To track the *Cac*/*Clj* species ratio in the fermentations, RNA-FISH was employed as described using the ClosAcet and ClosLjun probes (Hill and Papoutsakis 2024). Briefly, 1 mL of media from each timepoint was frozen at -20°C. Later, the samples were thawed and combined with an equal volume of ice-cold 1xPBS, washed twice, and resuspended in a blend of 1:1 PBS and ethanol. The OD_600_ values of these samples were noted. Next, an OD_eq_ of 0.15 of each sample was pelleted and dried at 46°C to allow any ethanol to evaporate. The pellets were then mixed with 75 μL of a hybridization buffer containing, per 1 mL, 222 μL formamide, 554 μL DI water, 200 μL 5 M NaCl, 22 μL 1M Tris-HCl, 1.1 μL 10% sodium dodecyl sulfate, and the ClosLjun and ClosAcet probes. The samples were incubated for 5 hr at 46°C.

Next, the cells are pelleted at 10,000 rpm for 10 min. 500 μL of 48°C washing buffer containing, per mL, 926 μL DI water, 43 μL 5 M NaCl, 20 μL 1 M Tris-HCl, 10 μL 5 M EDTA, and 1 μL 10% sodium dodecyl sulfate, was added and allowed to incubate for 25 min at 48°C. This washing step was repeated once before the samples were pelleted and resuspended with 500 μL of 1xPBS. The samples were then refrigerated before being analyzed on a flow cytometer (CytoFLEX, Beckman Coulter) as described (Hill and Papoutsakis 2024).

### Metabolic Flux Analysis

The methodology of metabolic flux analysis used in this work is similar to that used previously (Papoutsakis 1984, Desai, Nielsen et al. 1999). The chemical species and metabolic reactions (depicted in Figure 1) for the stochiometric model used to generate the stochiometric coefficient matrix are described in Appendix 1. In order to apply this method to a co-culture containing two distinct species we adopted a *structured*, *unsegregated* modelling approach. In other words, the intracellular metabolites are treated as mathematically unique chemical species despite being chemically identical. For instance, acetyl-CoA generated from glucose in *Cac* (acetyl-CoA-Cac) and acetyl-CoA generated from the WLP in *Clj* (acetyl-CoA-Clj) are chemically identical but can only be used in metabolic reactions related to the organism within which it was generated. The particular electron co-factors (i.e. NAD and ferredoxin) used in *Cac* metabolism is based on what is currently known about electron pathways in *Cac* (Foulquier, Rivière et al. 2022). A non-specific electron carrier (i.e. ‘EC’) was used in the *Clj* reactions, because the electron pathways are less confidently known in *Clj*. This is an appropriate solution in the case of our model because we are not attempting to calculate ATP generation which is dependent on the interconversion of specific electron carriers (Katsyv and Müller 2020). Some assumptions were made to prevent the model from being underdefined. First, it was assumed that all fructose is consumed by *Clj*, despite *Cac*’s ability to consume fructose. This assumption is sound because we have consistently observed that the rate of fructose consumption in monocultures of *Cac* is significantly slower than in co-culture. Second, we assumed that *Clj*’s only metabolic product is acetate. ^13^C-MFA on mixotrophically grown *Clj* shows that acetate is the sole product of energy metabolism (Dahle, Papoutsakis et al. 2022). The code used to solve the matrix equations for each data set was written in Python with Numerical Python and SciPy for matrix solving functions. Spyder was used as the integrated development environment. The code was compiled and tested on a Dell Inspiron 7390 with an Intel Core i7-8565U CPU and 16 GB of RAM.

**Figure 1:**
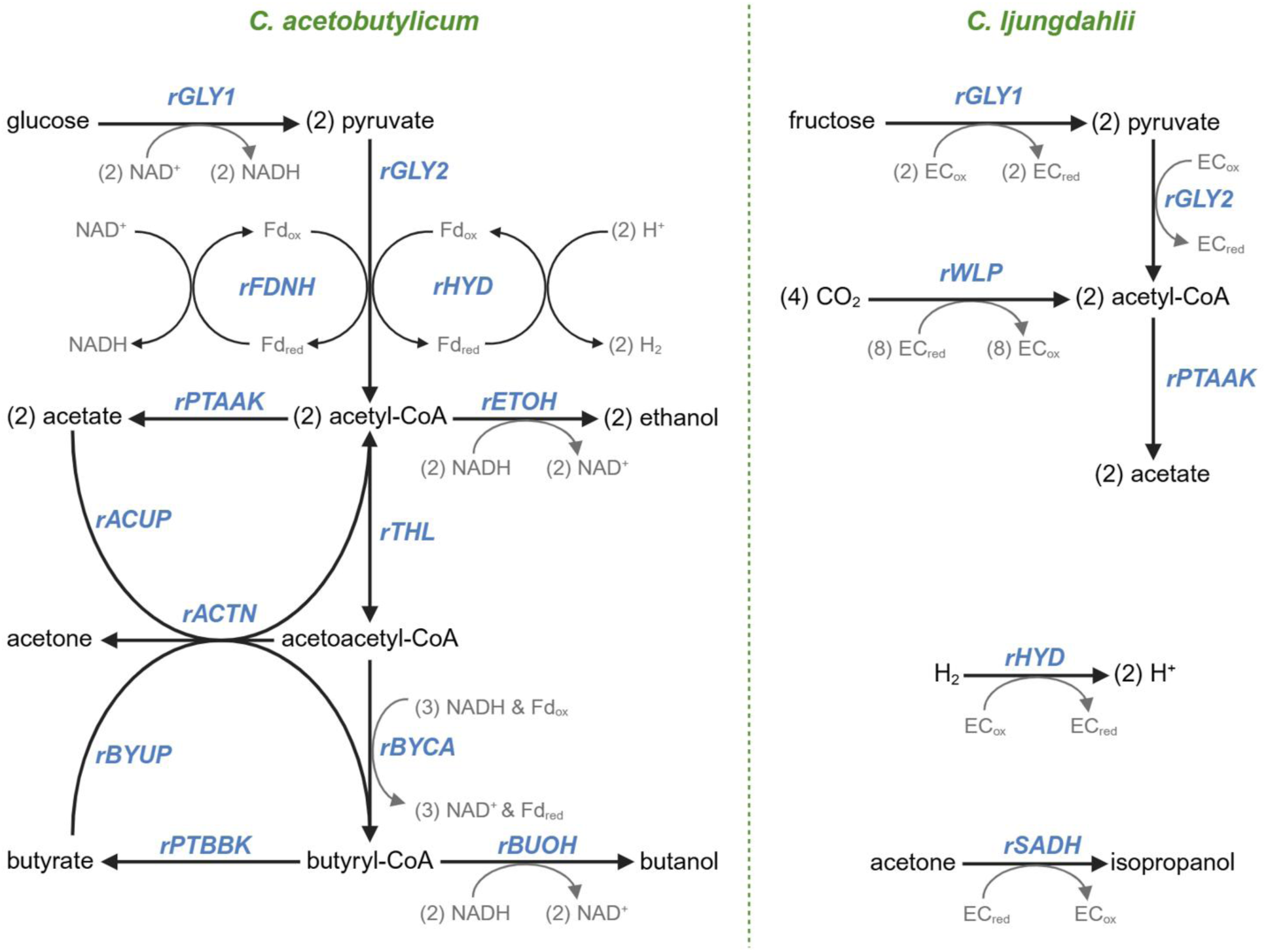
A metabolic map and associated fluxes of the primary metabolism of *Cac* (left) and *Clj* (right). In *Cac*, glycolysis catabolizes glucose to pyruvate (the rGLY1 flux). Pyruvate decarboxylation (rGLY2) forms acetyl-CoA, with electrons being conserved by forming (rFDNH flux) reduced ferredoxin (Fd_red_) which facilitates the conversion of protons to H_2_ (rHYD). Acetate (rPTAAK) and ethanol (rETOH) are produced from acetyl-CoA. The thiolase responsible for rTHL flux that catalyzes the formation of acetoacetyl-CoA from acetyl-CoA is expressed from the corresponding chromosomal gene (*thl*) and also from the copies of the same gene on the p95ace02a plasmid. Acetate (rACUP) and butyrate (rBYUP) reuptake via the CoA transferase is coupled to acetone formation (rACTN) from acetoacetyl-CoA. The three enzymatic step conversion of acetoacetyl-CoA to butyryl-CoA corresponds to the rBYCA flux. From butyryl-CoA, butyrate production is carried by the rPTBBK flux, and butanol production by the rBUOH flux. In *Clj*, fructose is catabolized to acetyl-CoA through the corresponding rGLY1 and rGLY2 fluxes. Acetyl-CoA is synthesized by the WLP pathway (the rWLP flux) from CO_2_. Acetate is synthesized through the rPTAAK pathway. *Clj* takes up (rHYD) H_2_ through its hydrogenase system and converts acetone to isopropanol (rSADH) using a seconday dehydrogenase. A non-specific electron carrier (‘EC’) is used in the *Clj* reactions because either several electron carriers exist or the electron carriers are not confidently known.

## Results and Discussion

### 1. *Cac*-ace02a and *Clj* cocultures in serum bottles with a H_2_ and CO_2_ atmosphere achieve superior product yields

First, we tested the impact of an H_2_ and H_2_/CO_2_ atmosphere on the *Cac*/*Clj* coculture. This phenomenon is especially critical in cocultures because of the acetogenic consumption of H_2_ and CO_2_ by *Clj* and the impact of H_2_ on *Cac* metabolism, whereby H_2_ removal at the local level due to utilization by *C. ljungdahlii* benefits *C. acetobutylicum* by reducing the feedback inhibition of its hydrogenase (Willis and Papoutsakis 2025). While pH-controlled bioreactors are preferred, we used 160-mL serum bottles (20 mL liquid volume) because pressure increases the H_2_ availability to *Clj* and due to safety concerns with continuous, pressurized H_2_ feeding into small-scale glass bioreactors at the university setting. Two biological replicates of four conditions (Figure 2) were tested: monocultures with an N_2_ headspace, cocultures with an N_2_ headspace, cocultures with an H_2_ headspace, and cocultures with a headspace of 80% H_2_ and 20% CO_2_. The pH in the serum bottles was manually controlled with two base additions at 10 and 27 hours (Figure 2H). It mostly remains in the range of 5.0-5.75.

**Figure 2:**
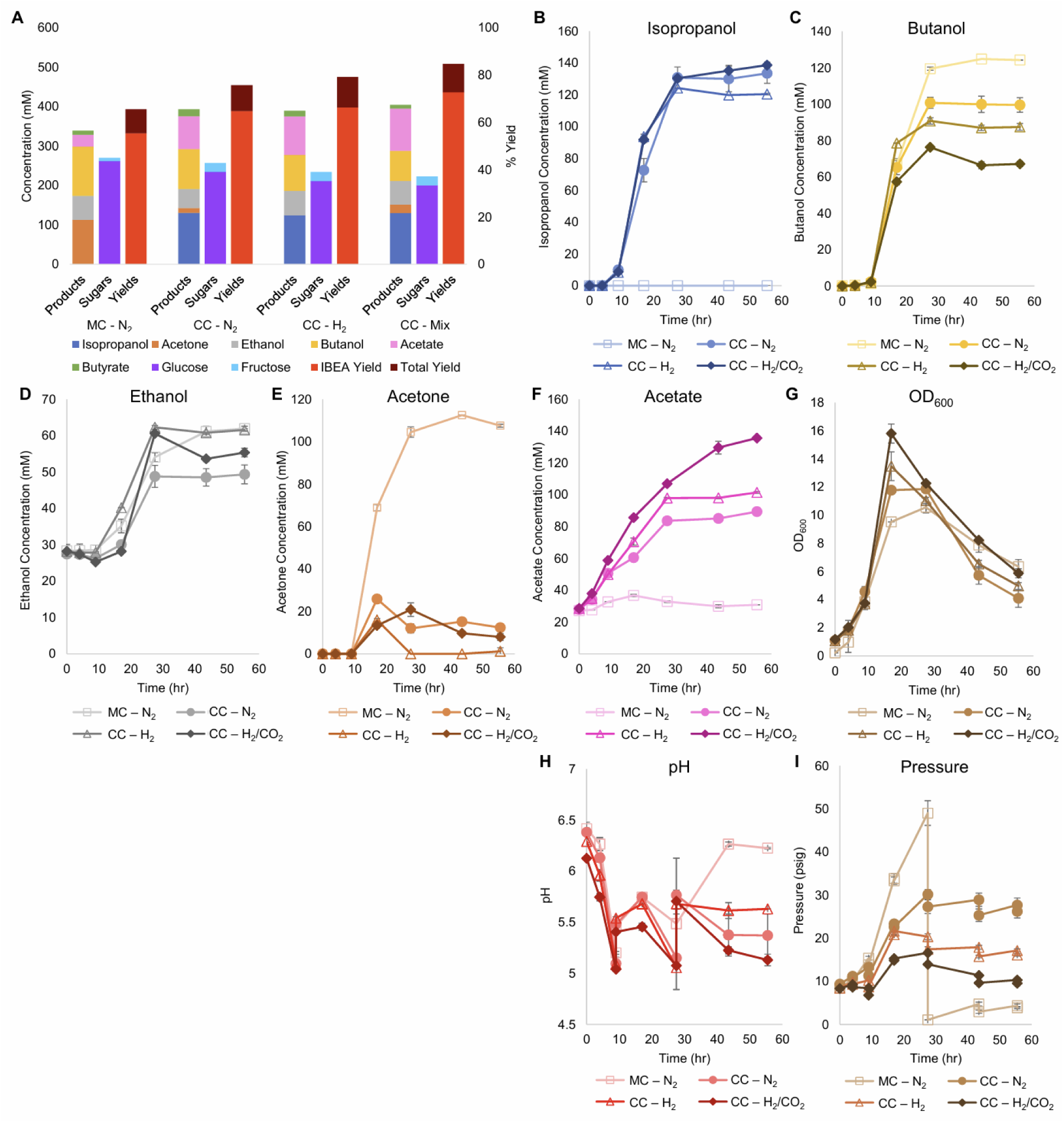
Metabolite production, sugar consumption, and metabolite yields of *Cac* and *Clj* cocultures in sealed serum bottles with different headspace gas composition: N_2_, H_2_ or Mix [H_2_/CO_2_ (80/20)]. MC refers to monocultures and CC refers to cocultures. (A) Summary of product concentrations, sugar consumption, and metabolite yields of the cocultures. The IBEA yield refers to IPA, BuOH, EtOH, and acetone. The total yield also includes butyrate and acetate. In all cases, the coculture yields surpassed the monoculture yields, primarily through the production of higher combined concentrations of acetone and IPA versus acetone alone in the monoculture, and through the production of additional acetate. (B) Kinetic profile of IPA formation. The monoculture produces no IPA. (C) Kinetic profile of BuOH formation. The monoculture produced a statistically significantly higher amount of BuOH than any coculture. (D) Kinetic profile of EtOH formation. (E) Kinetic profile of acetone formation. In the cocultures, acetone accumulation is typically less than 20 mM. (F) Kinetic profile of acetate formation. In contrast to the CO_2_-containing cocultures, the monoculture produced barely any acetate. (G) Biomass (OD_600_) formation kinetics. The cocultures reached the highest peak OD_600_. (H) The pH kinetic profile. The pH of the bottles was manually adjusted at 9 and 28 hours. (I) The pressure, in psig, of the sealed serum bottles. The monoculture bottles were bled of excess pressure at 28 hours to avoid explosion. H_2_-containing cocultures (including the mixed H_2_/CO_2_ gas) displayed the lowest unadjusted gas headspace levels, thus demonstrating the consumption of both gas components.

Given that glucose utilization generates large quantities of H_2_ and CO_2_ (Papoutsakis 1984) (Figure 1), with a N_2_ headspace in the coculture, pressures rose, indicating that more gas was produced than consumed by the cells. All bottles were initialized at 10 psig. The N_2_ monoculture was depressurized at 28 hours at 50 psig to prevent bottle rupture, but cumulatively, 40 psi of pressure was produced. With both H_2_ and H_2_/CO_2_, the headspace pressure increased minimally and eventually decreased, demonstrating significant gas reassimilation (Figure 2I). The H_2_/CO_2_gas coculture peaked at 16 psig before returning to 10 psig, the starting pressure, thus giving rise to virtually no pressure accumulation.

The presence of *Clj* resulted in increases in both the IBEA (Isopropanol, Butanol, Ethanol, Acetone) and total yields of this fermentation (Figure 2A), whereby the 80% H_2_ and 20% CO_2_ atmosphere performed the best. The IBEA (BEA correctly as no isopropanol is produced by WT *Cac*) yield for the monoculture is 55.4% with the total yield at 65.6% right below the theoretical maximum of 66.7% for a monoculture, as approximately a third of all sugar carbon is lost to CO_2_ (Papoutsakis 1984). The benefits of *Clj* in coculture can be seen beginning with the N_2_ headspace. While the BEA yield is 64.8%, acetate and butyrate production combine for an additional 10.9%, leading to a total yield of 75.7%. *Clj*, likely through direct cell-to-cell transfer (Charubin and Papoutsakis 2019, Charubin, Modla et al. 2020), is able to assimilate the CO_2_ produced by *Cac* and use the H_2_ electrons for additional energy generation. The total yield increased to 79.2% with H_2_ and 84.7% with 80% H_2_ and 20% CO_2_ (Figure 2A). Due to electron cofactor interconversion constraints, the yield for acetone production from glucose by *Cac* monocultures is described by Equation 1 below (Willis, Otten et al. 2025).

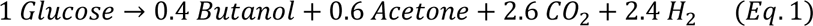

If both glucose and fructose are considered as sugar substrates, the 80% H_2_ and 20% CO_2_ bottles produced a yield of 0.68 mol 3C (IPA) per mol of sugar (Table S1), exceeding the monoculture yield of 0.42 mol 3C (acetone) per mol of sugar. If only glucose is considered, both the N_2_ and H_2_ bottles produced the theoretical maximum, and the 80% H_2_ and 20% CO_2_ bottles produced a yield of 0.76 (IPA). This result is consistent with our previous findings which show that *Clj* enables 3C yields in coculture that cannot be achieved in monoculture *Cac* fermentations (Willis, Otten et al. 2025).

All cocultures produced at least 130 mM of IPA (Figure 2B) with very little residual acetone (<20 mM), but the *Cac* monocultures produced about 110 mM of acetone (Figure 2E). In coculture, the presence of *Clj* led to production of more 3C metabolites (IPA and acetone): all cocultures produced at least 130 mM of IPA (Figure 2B) with very little residual acetone, but the monocultures produced about 110 mM of acetone (Figure 2E). This shows that the presence of *Clj* “coaxes” more acetone production by *Cac*, consistent with our previous observations (Charubin and Papoutsakis 2019, Willis, Otten et al. 2025). This was accompanied by reduced BuOH production, from ∼125 mM (in monoculture) to ∼65 mM in the coculture with H_2_/CO_2_ (Figure 2C). Furthermore, the presence of *Clj* resulted in significant increases in the total cell mass (OD_600_) in the fermentation, with the H_2_/CO_2_ conditions being the highest: an OD_600_ difference of about 5.5-6 units within about 15 hours (Figure 2G). Such OD_600_ levels cannot be accounted by *Clj* growth alone as *Clj* does not grow to high cell densities on H_2_/CO_2_, which are typically less than 2.5-3 OD_600_ units (Willis and Papoutsakis 2025). This suggests that *Clj* enhances the growth of *Cac* likely due to H_2_ consumption (Willis and Papoutsakis 2025) and through direct cell-to-cell contact (Charubin and Papoutsakis 2019, Charubin, Modla et al. 2020).

Significant differences in core metabolic fluxes between the culture conditions are depicted in Figure 3, with the remaining fluxes being largely the same. One difference between monoculture and cocultures is the distribution of carbon fluxes (rTHL, rBYCA, rACTN) through the acetoacetyl-CoA node in *Cac*. Before 10 hours, virtually all acetoacetyl-CoA is fed into the rBYCA pathway (Figures 3D & G), and very little acetone (rACTN) is produced (Figure 3H). In the cocultures, about half of carbon is directed through the rACTN reaction (Figure 3H). The early induction of rACTN pathway is probably a response to rising acetate concentration in the cocultures, because the flux through rACUP is high during this time, but rBYUP is nearly zero (Figure 3B). The butyrate uptake pathway becomes active around 12 hours. *Clj* is capable of activating the rACUP reaction since there is negligible flux through rACUP in the monoculture. This is a significant and unanticipated finding. We interpret this to be the result of higher rates of acetate production early in culture due to the growth associated with acetate formation by *Clj* (Figure 2F). In *Clj*, higher WLP fluxes in cocultures with H_2_/CO_2_ were expected because of the theoretically higher availability of CO_2_ and H_2_ to support autotrophic growth (Figure 3L). The resulting higher acetate production may explain why the rACUP flux stayed higher later into the culture in cocultures with H_2_/CO_2_, compared to cocultures with N_2_ or H_2_. This is more evident between 10 and 20 hours. This is also mirrored in the higher rPTAAK and rHYD fluxes (Figures 3J-K).

**Figure 3:**
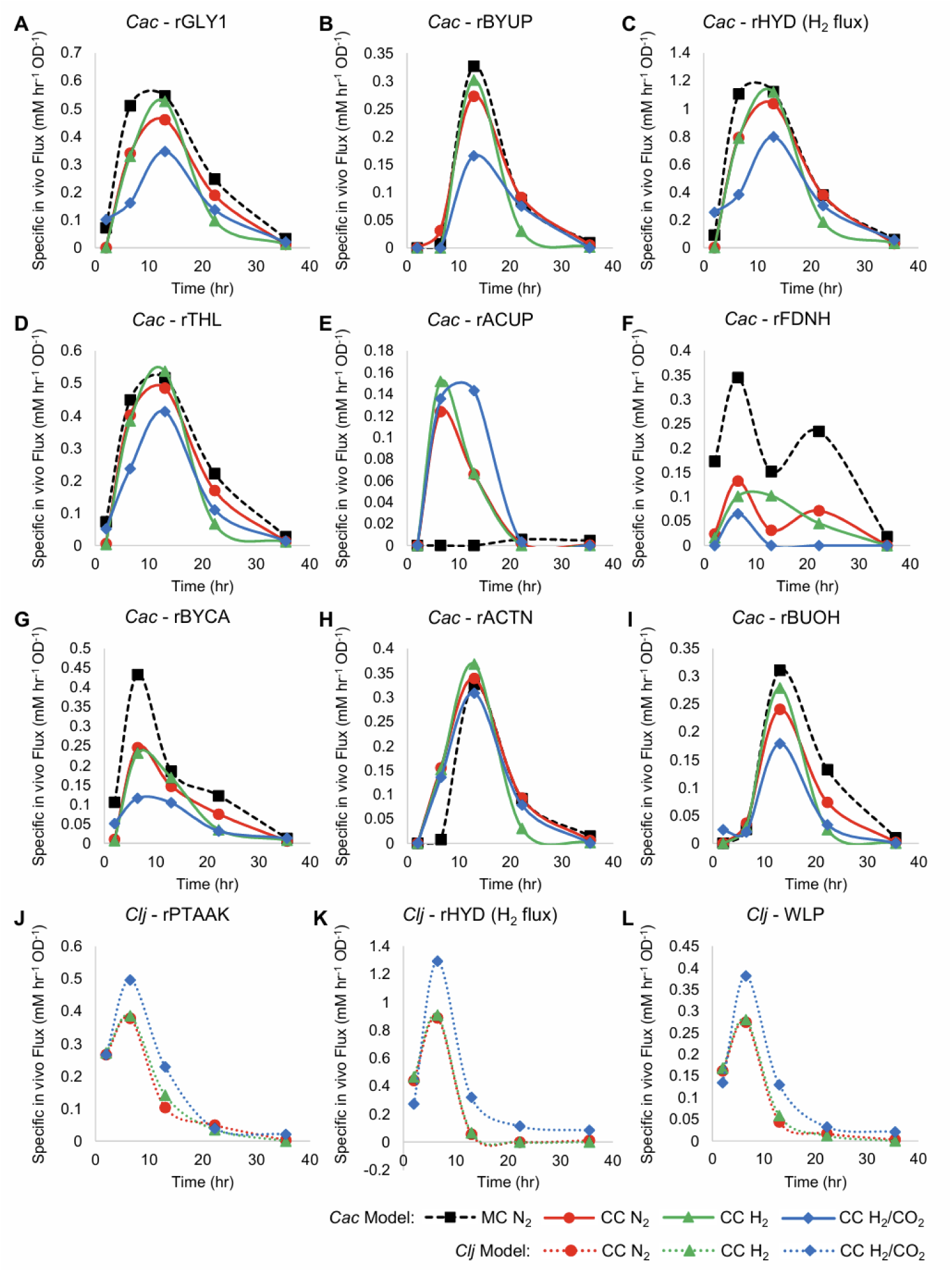
Fluxes of the monocultures (MC) and cocultures (CC) grown in pressurized serum bottles in units of mM/h/OD_600_, for the data of Figure 2. The flux maps are shown in Figure 1. *Cac* fluxes are depicted in panels A-I, and *Clj* fluxes are depicted in panels J-L with dotted lines. The *Cac* monoculture with a nitrogen headspace is depicted with thick black dashed lines. The *Cac* monoculture has notably higher rGLY1, rBYUP, rHYD, rFDNH, rBYCA, and rBUOH fluxes. However, unlike the cocultures, it displays to no acetate uptake (E) (rACUP). Compared to the monoculture, in cocultures with the H_2_/CO_2_ atmosphere, *Cac* shows lower fluxes for rGLY1, rBYUP, rHYD, rTHL, rBYCA, and rBUOH, but higher fluxes for acetate uptake (rACUP). *Clj* clearly has higher fluxes with H_2_/CO_2_ atmosphere, suggesting the benefits of CO_2_ availability in the coculture setting. The pure N_2_ and pure H_2_ atmospheres performed similarly, suggesting that the use of CO_2_ generated by *Cac* did not require additional, exogenous H_2_ beyond the H_2_ produced by *Cac*.

In the *Cac* monoculture, a higher percentage of excess Fd_red_ is used to generate NADH than in the cocultures, especially after 10 hours (Figure 3F). As expected, this is correlated with butanol production. Butanol production requires NADH, which can be generated via the rFDNH reaction from excess Fd_red_. At the same time, the higher production of butyryl-CoA in the *Cac* monoculture, an intermediate in the butanol production pathway, produces additional Fd_red_ (Figure 1). The net result is a higher flux through the rFDNH reaction. In the coculture, butanol production is less strong (Figure 2C), probably resulting in lower Fd_red_ production. In summary, the cocultures result in better carbon management as well as improved solvent yields and selectivity (Figure 2A). In order to address two key deficiencies, lack of good pH control (Figure 2H) and incomplete utilization of glucose (Figure S4B), we next utilized controlled-pH bioreactors.

### 2. Exploring core process parameters (pH, intermittent *Clj* additions, and gassing configurations) for better coculture performance

#### 2.1 Evaluating the role of pH setpoints and intermittent *Clj* additions in coculture performance

With eight bioreactors, we tested a matrix of pH setpoints and intermittent *Clj* additions to assess their impact on metabolite yields. We considered the following: pH setpoints of 5.5 and 5.9, if a drop to pH 5.0 by 15 hours is beneficial, and if additional *Clj* would be needed for full acetone conversion or additional acetate production. Past research has indicated that a drop in pH to 5.0 may be necessary to facilitate a shift in *Cac* from acidogenesis to solventogenesis (Grupe and Gottschalk 1992, Girbal, Croux et al. 1995, Grimmler, Janssen et al. 2011, Haus, Jabbari et al. 2011). To allow for this drop, the one-way pH controllers were set to maintain a floor of 5.0 after the bioreactors were initialized at a pH of 5.5 or 5.9. The acids produced by the cells would lower the pH until the controllers initialized pH control at 5.0. The addition of *Clj* was proposed both to allow for more complete acetone conversion – if needed – and to replenish the *Clj* population if it was negatively impacted by the low-pH or pH-recovery period. For the bioreactors that received additional *Clj*, 160 mL of *Clj* at an OD_600_ of 0.7 was concentrated and added to the bioreactors. Our results showed higher yields and increased BuOH and EtOH production with a pH of 5.5 (Figures S1 A-B, E-F; S2 G-H). Isopropanol concentrations (titers) were similar across all conditions (Figures S1 C-D), and no significant acetone accumulation was observed with intermittent addition of *Clj*. At a pH of 5.9, higher concentrations of acetate and butyrate were produced than with a pH of 5.5 (Figures S2 C-F). Based on these results, we thus concluded that both the additional bolus of *Clj* when using these bioreactors without pressurized H_2_/CO_2_ (with the proviso that this may not hold when H_2_/CO_2_ are supplied at high pressure) and a pH drop to 5.0 were not needed. Notably, glucose consumption was higher in the bioreactors compared to the serum bottles due to the stable control of pH. We chose to move forward with a pH of 5.5 to produce a greater balance of IBEA and lower concentrations of acetate and butyrate.

#### 2.2 Evaluating different bioreactor gassing configurations

While the serum bottles produced high yields of our target molecules, they lack online pH control and cannot be practically scaled up. Thus, we utilized biological replicates in bioreactors with three gassing conditions: no gassing, headspace gassing only, or sparging. In this coculture, *Cac* produces H_2_ and CO_2_, and *Clj* consumes these gases (Charubin and Papoutsakis 2019). Ideally, we would have liked to gas the bioreactors with exogenous H_2_/CO_2_ (80/20), but safety issues in these bioreactors, which lack sufficient exhaust in our labs, prevented that. We still wanted to examine the impact of two key modes of gassing and compare to no gassing as explained below. Thus, for all bioreactors, a blend of 85% N_2_, 10% CO_2_, and 5% H_2_ was selected at a flowrate of 10 mL/min. This allows for some addition of exogenous H_2_ and CO_2_ supply, while remaining below flammable limits. Two issues are associated with gassing from the physical point of view. We want to avoid any stripping of the IBEA solvents while also avoiding O_2_ intrusion. Sparging, or direct bubbling of the gas into the fermentation media, would strip more IBEA than any other option. Headspace gassing would be expected to lead to less IBEA stripping, but it is minimally effective in terms of gas transfer due to a very small interfacial mass transfer rate (k_L_a). No gassing, which is when the gassing was only enabled during sampling to account for the very minor decrease in media volume, would be associated with the least stripping IBEA while potentially allowing O_2_ intrusion. IBEA production and productivity of the coculture was similar across all gassing conditions (Figure S3). No negative impact on cell growth or IBEA production was observed with no gassing, suggesting that O_2_ intrusion is not a concern. Concentrations of BuOH and IPA were similar across all gassing conditions after 50 hours, indicating that there was no significant stripping of these metabolites. We thus conclude that any gassing configuration is suitable for future fermentation experiments.

### 3. Exploring impact of starting (inoculation) cell densities (SCD) and species population ratios in *Cac-Clj* cocultures and, for comparison, in *Cac* monocultures

Industrial fermentations are initiated with high cell densities in order to minimize the lag phase and shorten the duration of the fermentation. In this spirit, we next probed the role of different starting cell densities (SCDs) on metabolite production in cocultures and, for comparative reasons, in *Cac* monocultures, as *Cac* is the organism that metabolizes glucose and is responsible for BEA production. We also used RNA-FISH (Hill and Papoutsakis 2024) to evaluate the changes in cell population ratios over time, which is important for the industrial scaling up of coculture fermentations..

#### 3.1 A low starting cell density (SCD) is best for metabolite production by *Cac* monocultures

To goal of these experiments was to establish a baseline for *Cac*-ace02a (which has a different metabolite profile compared to the WT *Cac*) fermentation and examine whether higher SCDs of *Cac* would improve metabolite yields and titers. For ease of comparison with future coculture experiments, the bioreactors were sparged with 85% N_2_, 10% CO_2_, and 5% H_2_ at 5 CC/min. For the high SCD fermentations, the inocula were 3x as concentrated as the standard (referred to as low below) SCD fermentations, so the high SCD fermentations were inoculated at an OD_600_ of 0.3 while the low SCD fermentations were inoculated at 0.1. The low-SCD monocultures produced higher titers of BuOH (200 mM) and acetone (152 mM) while consuming less sugar (Figure 4). Thus, the yields in both low-SCD fermentations were higher. Similarly, the ratio of the carbon moles in fermentation products compared to the carbon moles of sugar consumed was 0.5 with a low SCD and 0.4 with a high SCD (Figure 4F).

**Figure 4:**
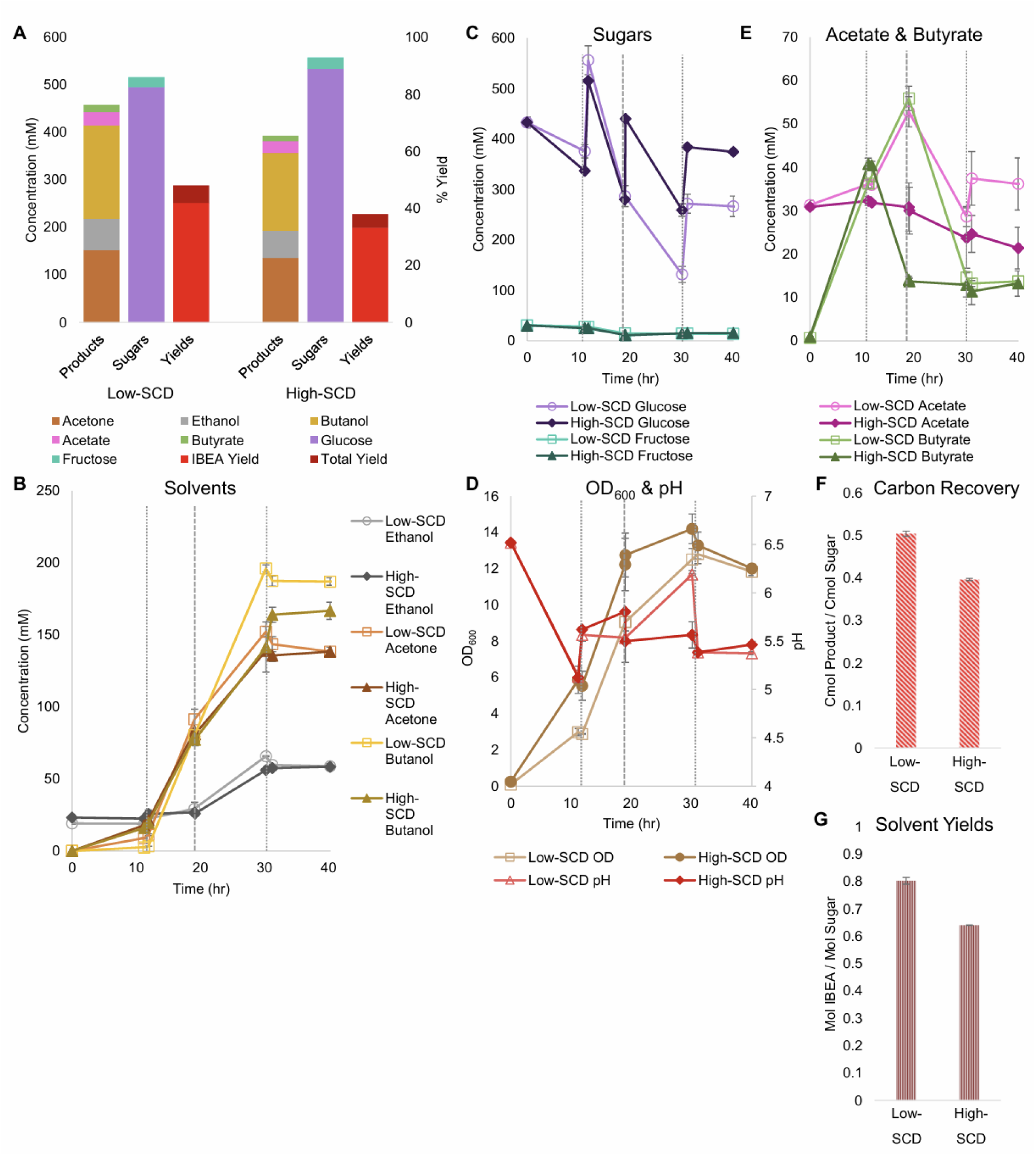
Metabolite production, sugar consumption, metabolite yields and carbon recoveries into metabolites of *Cac* monocultures in bioreactors at low- and high starting cell densities (SCDs). Glucose was fed thrice in the high-SCD operation, at 10, 20, and 30 hours, and twice in the low-SCD operation, at 10 and 30 hours. The vertical, short dashed lines reflect glucose additions to all bioreactors. The vertical, long dashed lines at 20 hrs reflect glucose additions to only the high-OD bioreactors. (A) Summary of sugar consumption, metabolite concentrations and yields. The IBEA yield refers to IPA, BuOH, EtOH, and acetone combined. Without *Clj*, the monoculture produces only acetone with no IPA. The total yield also includes butyrate and acetate. (B) The EtOH, acetone, and BuOH formation kinetics. (C) The kinetic profile of sugars in the bioreactors. (D) The OD_600_ (left y-axis) and pH (right y-axis) kinetic profiles. (E) The acetate and butyrate concentration kinetics. (F) Carbon-mol (C-mol) of products produced per C-mol of sugars consumed. (G) Mol of alcohols (IBEA) produced per mol of sugar consumed.

In terms of maximum productivity, the low-SCD condition produced BuOH at 13.7 mM/hr, EtOH at 4.4 mM/hr, and acetone at 9.1 mM/hr. The high-SCD condition produced BuOH at 11.1 mM/hr, EtOH at 3.8 mM/hr, and acetone at 9.2 mM/hr. While the *Cac* in the high-SCD bioreactors grew more quickly and to a slightly higher OD_600_, by 30 hours, and through the rest of the fermentation, the effects of the higher SCDs disappeared (Figure 4D). The pH values of the bioreactors were well-controlled at 5.5 after an initial drop to the controllers’ baseline and a manual addition of HCl at 30 hours. In the end, the low-SCD monocultures produced 0.8 mol of BEA per mol of sugar, and the high-SCD conditions produced 0.64 mol/mol (Figure 4G).

For monoculture fluxes, the primary differences between the metabolism of cells when inoculated at high or low SCDs are the higher specific fluxes seen in the low-SCD condition prior to 24 hours (Figure 5). After 24 hours, the fluxes are almost indistinguishable, with the exception of the acetate uptake reaction (rACUP), which suggests a second, late phase of acetate uptake (Figure 4E). Overall, at high SCDs, the lower acetate concentrations (Figure 4E) are a reflection of higher rates of acetate uptake (rACUP; Figure 5E). The lower butyrate concentrations (Figure 4E) reflect better flux management around the acetyl-CoA node (Figures 5A, D, & G). The lower butyrate formation at higher SCDs results in lower butanol formation and higher acetone selectivity [ratio of acetone to (butanol + ethanol)]. This behavior is also correlated with a lower efficiency of sugar utilization at higher SCDs, as evidenced by the carbon and solvent yields (Figure 4F-G).

**Figure 5:**
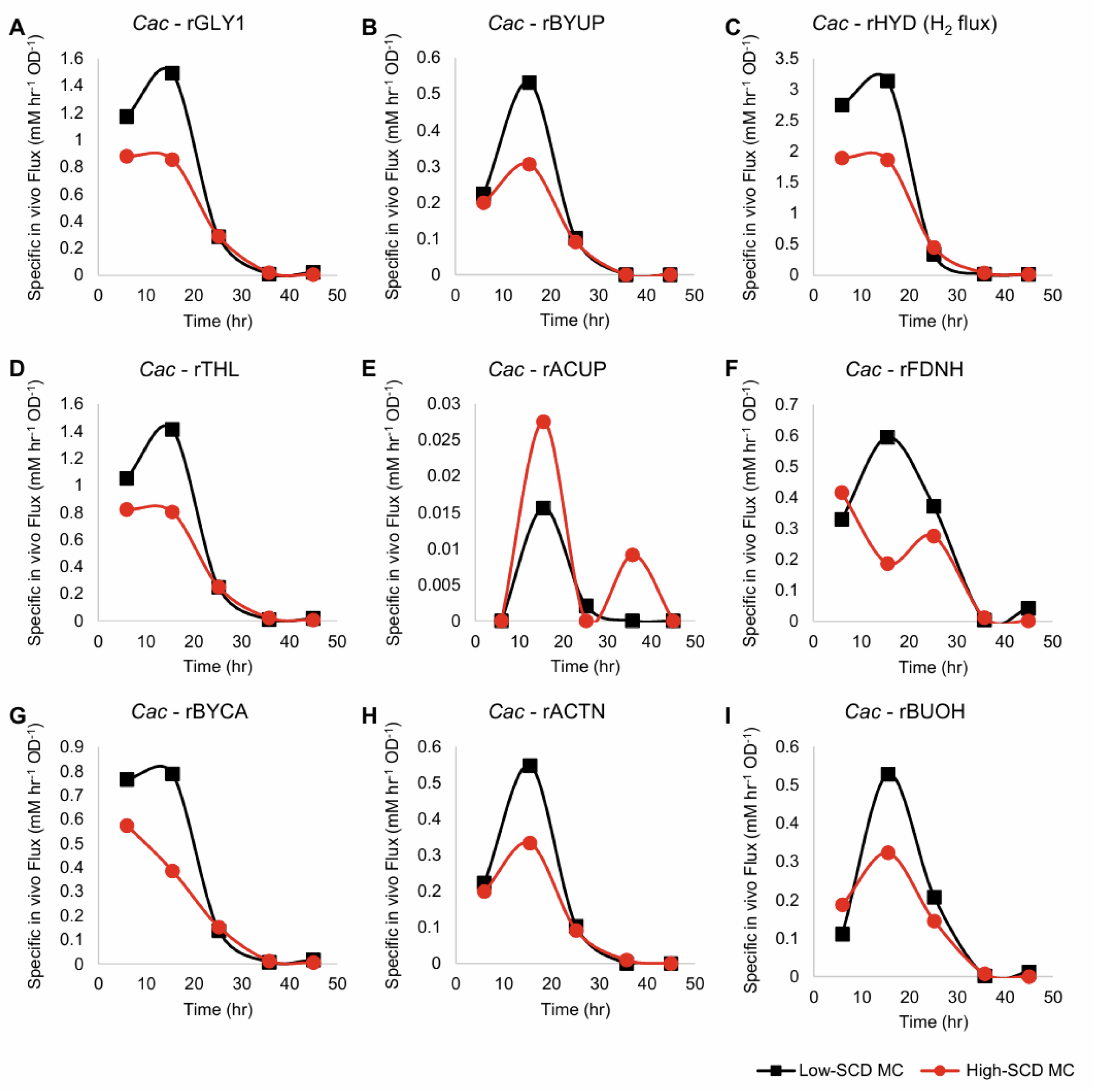
Metabolic fluxes (mM/h/OD_600_) of the monocultures corresponding to Figure 4. With the exception of rACUP, the cultures with a lower SCD displayed higher fluxes.

#### 3.2 Higher coculture starting cell densities (SCDs) result in higher metabolite yields and shift the product balance towards isopropanol

For the experiments to assess the impact of higher SCDs in *Cac*/*Clj* cocultures, all experimental conditions were identical to those in section 3.1 except for the addition of *Clj* at the beginning of the fermentation. The low SCD fermentations were inoculated with an OD_600_ of 1 and the high SCD fermentations were inoculated with an OD_600_ of 3 with the same species ratio of 1 *Cac* : 8 *Clj.* To gain clarity about the ratio of each species’ cell population during the fermentation, RNA-FISH was performed across all timepoints (Figures 6D&E; S5D&E). Our RNA-FISH method (Hill and Papoutsakis 2024) identifies the viable population of each species. The non-viable cells are not stained or labeled and are displayed as “gray” in Figures 6D&E. RNA-FISH analysis showed that *Clj* predominated, by design, early in the cocultures, but by 10 hours, *Cac* comprised the majority of cells. The high-SCD condition maintained a larger fraction of live *Clj* cells for longer, although the low-SCD condition ended with more live cells of both species.

**Figure 6:**
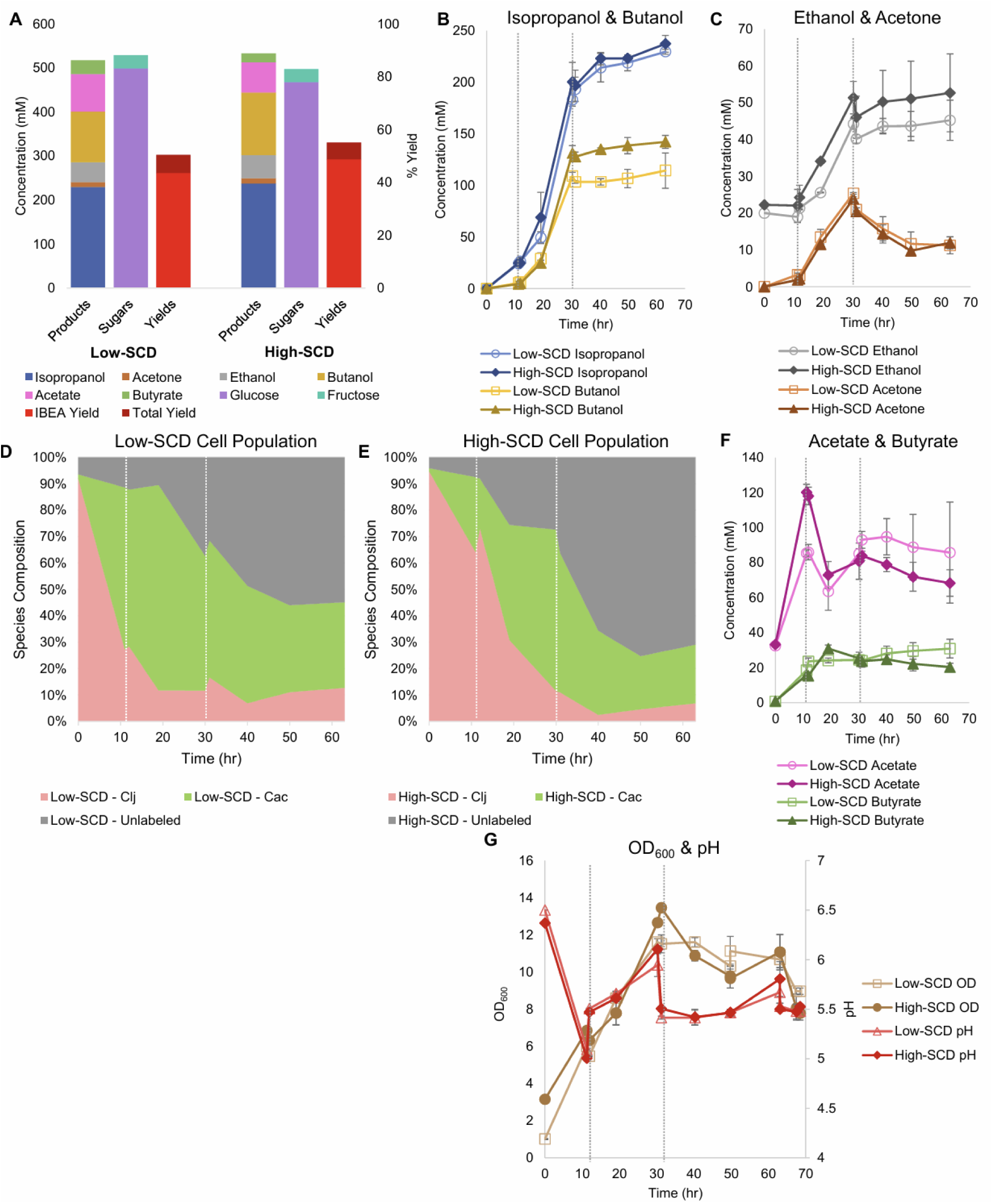
Sugar consumption, metabolite production and yields, and total biomass (OD_600_) of low- and high-SCD *Cac* and *Clj* cocultures. The vertical, short-dashed lines represent glucose additions to the bioreactors. (A) Summary of sugar consumption, metabolite concentrations and yields. IBEA yield refers to only IPA, BuOH, EtOH, and acetone. The total yield also includes butyrate and acetate. (B) IPA and BuOH formation kinetics. (C) The EtOH and acetone formation kinetics. (D-E) Species composition over time, as determined by RNA-FISH labeling for (D) the low-SCD and (E) the high-SCD. (F) The acetate and butyrate concentration kinetics. *Cac* can produce and uptake both acetate and butyrate; acetate is the main product of *Clj*. (G) The OD_600_ (left y-axis) and pH (right y-axis) kinetic profiles.

IPA (230-237 mM), rather than BuOH (114-142 mM) or acetone (final concentrations of 12 mM), was the dominant product, and the IBEA yield exceeded 48% in the high-SCD condition (Figure 6A). The total yield was 55%. Nearly 0.56 carbon moles of metabolites were produced for every carbon mole of sugar consumed in the high-SCD condition (Figure S5B). In one high-SCD replicate, 0.93 moles of IBEA were produced for every mole of sugar consumed for an average of 0.89 mol/mol (Figure S5C). In the low-SCD condition, 0.77 mol IBEA/mol sugar was produced. In terms of maximum productivity, both conditions produced IPA at 13.9 mM/hr, which is markedly higher than the monoculture acetone production of 9.1-9.2 mM/hr. This again shows that the presence of *Clj* leads to production of more 3C solvents, suggesting that *Clj* alters *Cac*’s metabolism to benefit IPA/3C product formation. The low-SCD condition produced BuOH at 8.3 mM/hr, EtOH at 3.0 mM/hr, and acetone at 1.6 mM/hr. The high-SCD condition produced BuOH at 10.4 mM/hr, EtOH at 2.8 mM/hr, and acetone at 1.5 mM/hr. In terms of three-carbon yields, the high-SCD cocultures doubled the yields of the high-SCD monocultures, 0.25 to 0.53 when only glucose is considered as the sugar substrate (Eqn. 1, Table S1). The advantages of cocultures with CO_2_-capturing *Clj* in yields and product ratios are clear, and the shift in the product ratio to IPA from BuOH is notable.

Flux calculations corresponding to these cocultures (Figure 7) reveal that the uptake of carboxylic acids for acetone production displays two stages. Before about 20 hours, acetate but not butyrate is taken up (Figures 7B & E). After 20 hours, mostly butyrate is taken up. This results in a wide peak for the rACTN flux profile (Figure 7H), though the carboxylic acid used to receive the CoA for the CoA-transferase (CtfAB) reaction differs over time. Peak specific H_2_ production (rHYD) occurred earlier in the low SCD conditions (Figure 7C), and this is likely caused by the increased specific glycolytic rate compared to the high SCD condition. This mirrors the behavior observed in the *C. acetobutylicum* monocultures where the specific glycolytic rates were higher in the low-SCD condition.

**Figure 7:**
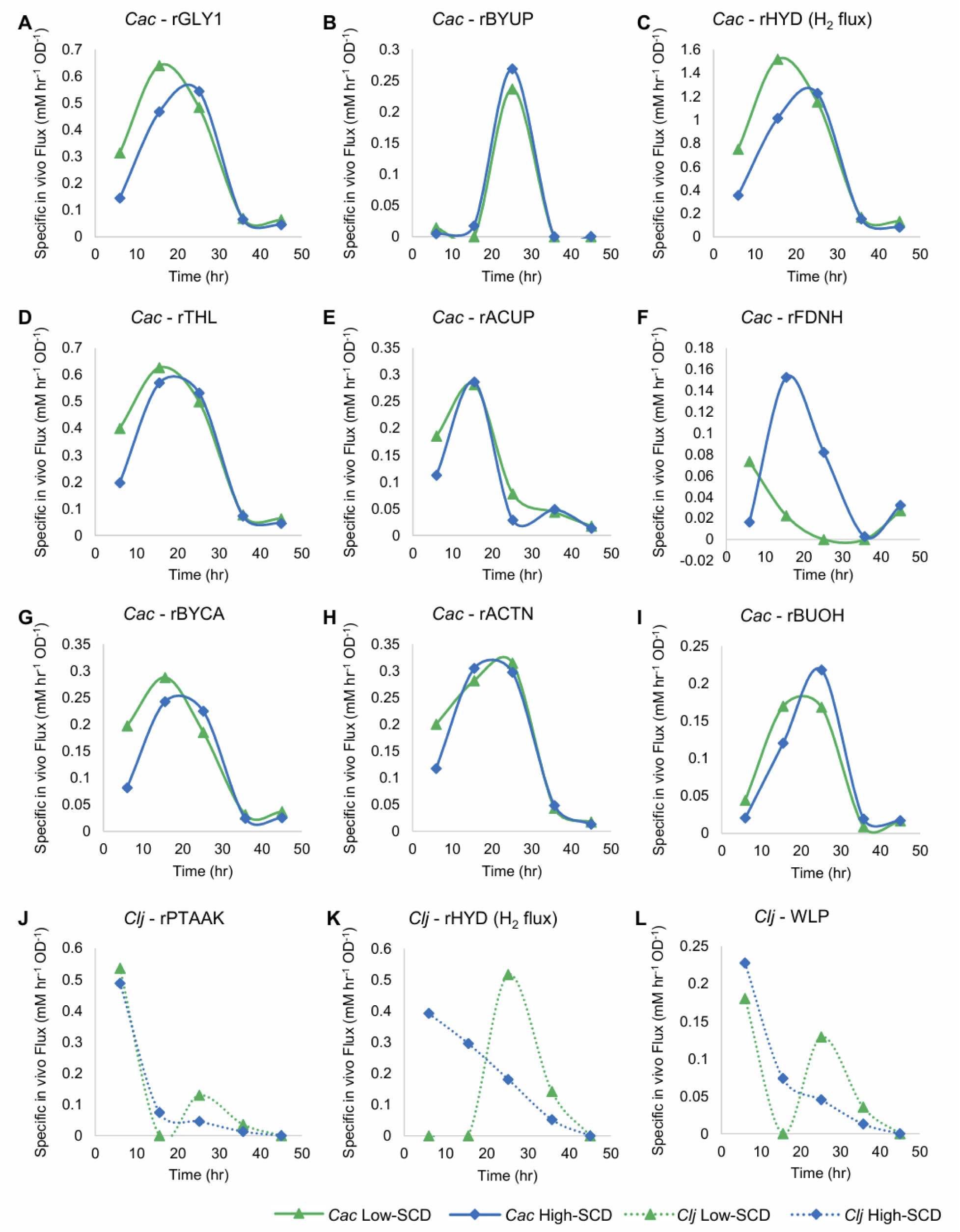
Metabolic fluxes (mM/h/OD_600_) of the cocultures shown in Figure 6. *Cac* fluxes are depicted with solid lines on panels A-I and *Clj* fluxes are depicted with dotted lines on panels J-L.

The difference in the rFDNH flux (NADH generation from Fd_red_) is very distinct between the two culture conditions: it starts at a modest level and decreases at the low SCD condition to reflect the need for reduced Fd to produce H_2_ earlier and at a higher specific rate compared to the high SCD culture. In the latter, the rFDNH flux started low but peaked to higher levels around hour 15 (to feed the rBuOH and other alcohol fluxes (not shown)), reflecting a delayed and lower H_2_ production (compared to the low SCD culture).

The differences between *Clj* activity in the high and low SCD conditions is stark. Hydrogen uptake (the rHYD flux) is very distinct, as it starts high and decreases monotonically in the high SCD condition, while remaining zero up to 15 hours and then increasing dramatically to reach a peak around 25 hours. This means that the high SCD cultures are electron-limited until the very end of the active cell growth period. In contrast, the high SCD cultures are electron-sufficient until about 15 hours becoming electron limited after that until the end of the active cell growth. What is also interesting is the period between 10 and 30 hours where, in *Cac*, rACTN was very high (Fig. 7H), consequently forming more of CO_2_ and H_2_ (compared to alcohol production) for *Clj*, but only at low SCD *Clj* seems to strongly engage its WLP during this time period as demonstrated by an increasing H_2_ uptake rate (rHYD) reaching a peak around 30 hours (Figure 7K).

These cocultures ultimately represent the highest-performing IB or IBEA fermentations to date. With IPA concentrations of nearly 250 mM, while maintaining strong BuOH production of 150 mM, these fermentations provide a strong possibility of sustainable biofuel production. Compared to prior literature, our result of 0.58 cmol product/cmol sugar is near the theoretical maximum of 0.65 (Charubin and Papoutsakis 2019). Note also that this past work, which only produced 193 mM of IPA, was done in serum bottles, which can be pressurized and do not allow for any solvent evaporation.

### 4. Comparison to prior literature demonstrates superior IBEA concentrations and improved productivities

Monocultures and cocultures have long been used to produce IPA and BuOH, often in blends and sometimes with acetone and EtOH as minor components as well. Our lab has discovered the expanded metabolic space that is enabled in cocultures of *Cac* and *Clj* (Charubin and Papoutsakis 2019) and further probed these interactions to discover the exchange of cytoplasmic proteins, RNA, and DNA (Charubin, Modla et al. 2020, Charubin, Hill et al. 2024). Our work, when contextualized in the literature, demonstrated higher IPA and BuOH titers and productivities (Figure 8, A vs B) than most similar studies. Our process produces mostly IPA on a molar basis. Crucially, our process is fast and efficient, producing most of the IBEA by 32 hours and finishing by 63 hours. Other cell systems reported fermentations lasting from 72 to 158 hours, leading to a productivity decline of more than half (Figure 8, Table 1). Systems based on engineered *Escherichia coli* (Jojima, Inui et al. 2008) and *Cupriavidus necator* (Boy, Lesage et al. 2023) monocultures have similar IPA productivities, but they do not produce butanol so the total IBEA productivity and titers were much lower compared to our approach. Another study using a different *E. coli strain* in fed-batch fermentation achieved exceptional IPA productivity and titers (662 mM, 1986 mM-C, 2.05 mM-C per mM glucose) in 71 hours, significantly exceeding our IBEA titer over a similar timescale (Inokuma, Liao et al. 2010). This level of IPA production required nearly twice as much glucose consumption (967 mM, with associated supplementation every ∼7 hours) compared to our fermentation (496 mM combined glucose and fructose) which produced 237 mM IPA and 142 mM butanol (1279 mM-C, 2.58 mM-C per mM glucose). This 26% percent increase in mol IBEA per mol glucose yield of our fermentations, compared to the results from Inokuma, Liao et al., illustrates the value of our coculture approach which uses *Clj* to recapture meaningful quantities of CO_2_ from glycolysis and improve the carbon efficiency of glucose fermentation. Other monocultures and cocultures of *Clostridium beijerinckii* produced lower IBEA titer, lower IBEA productivity, or both (Moon, Han et al. 2015, Procentese, Raganati et al. 2018, Dalal, Das et al. 2019, Rochón, Cebreiros et al. 2019, Cui, He et al. 2020, Carrié, Velly et al. 2022, Hocq and Sauer 2022).

**Figure 8:**
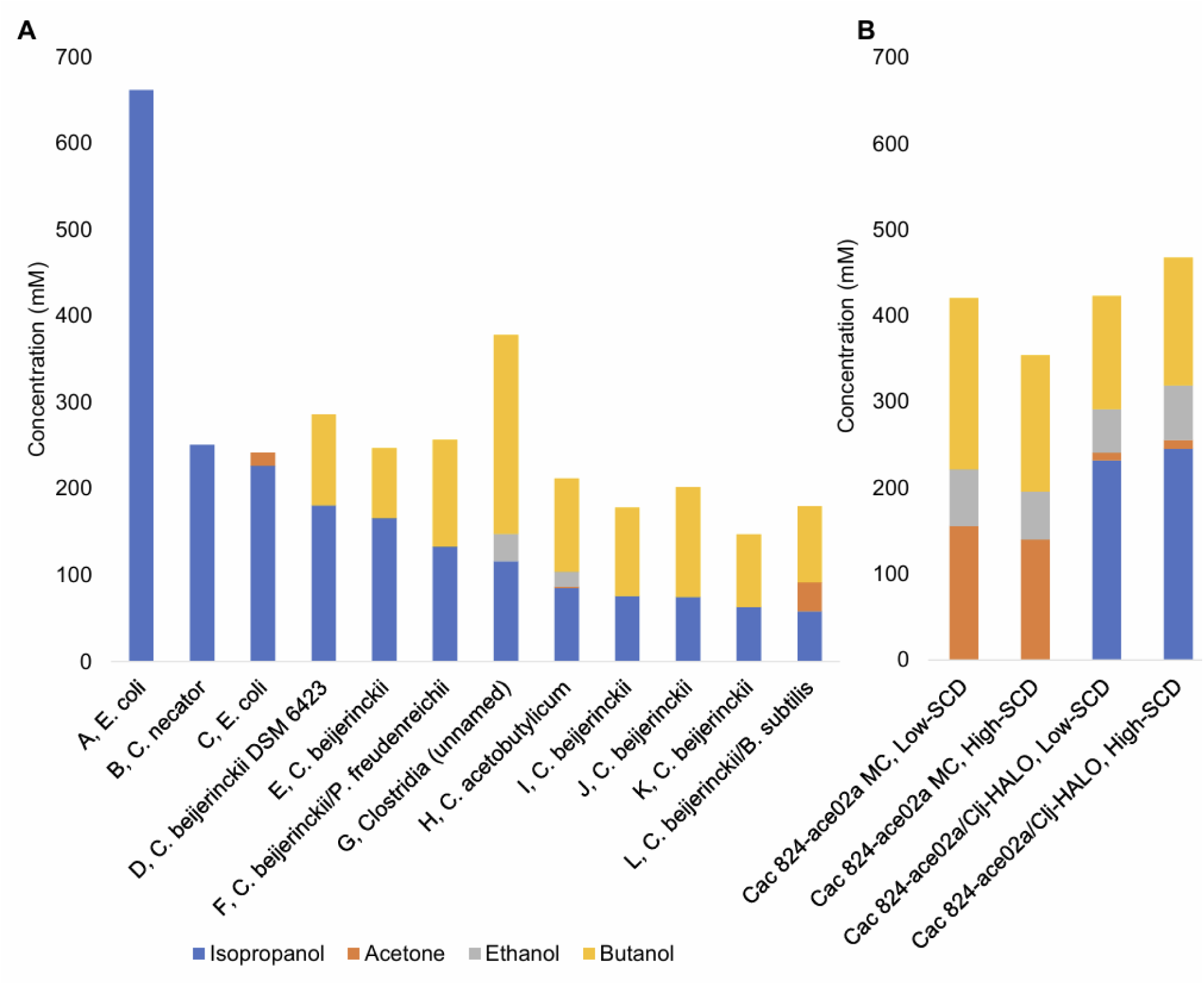
(A) Select metabolite concentrations in the literature for monoculture, coculture, and mixed culture productions of IBEA. The letters A-K preceding the commas correspond to the row headings in Table 1. (B) Data from this study for low-SCD and high-SCD *Cac* 824-ace02a monocultures and *Cac* 824-ace02a/*Clj*-HALO cocultures. Note the higher total IBEA titer and superior IPA:BuOH ratios.

**Table 1:** Select species used in the literature for monoculture, coculture, and mixed culture productions of IBEA. The letters A-L preceding the commas correspond to the bar labels in Figure 8.

| Letter | Species | Year Published | Time to Result (hrs) | Source in References |
| --- | --- | --- | --- | --- |
| A | *E. coli* | 2010 | 72 | (Inokuma, Liao et al. 2010) |
| B | *C. necator* | 2023 | 76 | (Boy, Lesage et al. 2023) |
| C | *E. coli* | 2008 | 36 | (Jojima, Inui et al. 2008) |
| D | *C. biejerrinkii* | 2019 | 158 | (Rochón, Cebreiros et al. 2019) |
| E | *C. biejerrinkii* | 2019 | 72 | (Dalal, Das et al. 2019) |
| F | *C. biejerrinkii* & *P. fruedenreichii* | 2022 | 120 | (Hocq and Sauer 2022) |
| G | *Clostridium* Str. WK | 2023 | 120 | (Zhang, Zhang et al. 2023) |
| H | *C. acetobutylicum* | 2012 | 48 | (Lee, Jang et al. 2012) |
| I | *C. biejerrinkii* | 2015 | 90 | (Moon, Han et al. 2015) |
| J | *C. biejerrinkii* | 2018 | 40 | (Procentese, Raganati et al. 2018) |
| K | *C. biejerrinkii* | 2022 | 50 | (Carrié, Velly et al. 2022) |
| L | *C. biejerrinkii* & *B. subtilis* | 2020 | 84 | (Cui, He et al. 2020) |

## Conclusions

There is a great need for renewable and sustainable production of essential chemicals from plant-based biomass, but with traditional fermentation techniques, a third of sugar carbon is lost as CO_2_. An effective way to overcome this limitation is through cocultures with an acetogen such as *Clj*. Through coculture cell interactions, we can both expand the metabolic space, enabling the production of isopropanol, while also capturing and reassimilating this CO_2_, bringing our yields to near-theoretical levels. Fortunately, this process is also efficient and scalable, with high isopropanol and butanol productivities that compare well to other proposed biological alternatives. Through RNA-FISH cell population tracking and coculture flux analysis, we have been able to build a superior model of coculture dynamics, observing significant differences in the rHYD and rFDNH fluxes across different experimental conditions. This work provides a significant foundation for further improvements. Genetic engineering tools for *Cac* and *Clj* are well-understood, so changes to their cell metabolisms and to the balance of fermentation products are possible. The coculture could also be enhanced with another member species, such as *Clostridium kluyveri*, or with expression of its genes, to enable chain elongation to produce 6-8 carbon products such as hexanol. Finally, from a process engineering perspective, continuous cultures, cell retention, and the feeding of H_2_ and CO_2_ could be implemented as this process is scaled to sustainably supply industrial chemical demands.

## Supplemental Material

**Supplemental Figures** (Figures S1-S5)

**Supplemental Table** (Table S1: “Calculated acetone/IPA yields with all sugars consumed (glucose and fructose) or with glucose only for each experimental condition.”)

**Appendix 1** (PDF) (“Stoichiometric Coefficient Calculations and Details”)

## Acknowledgements

This work was supported by an ARPA-E project under contract AR0001505. N.B.W. and J.D.H. were supported in part by a U.S. Department of Education GAANN Fellowship under grant P200A210065.

## Author Contributions

J.K.O., N.B.W., and E.T.P. designed research; J.K.O., N.B.W., A.D., and J.D. performed research; J.K.O., J.D.H., N.B.W., A.D., J.D. and E.T.P. analyzed data; E.T.P. supervised the project; and J.K.O., J.D.H., and N.B.W. wrote the paper.

---

**Table S1:** Calculated acetone/IPA yields with all sugars consumed (glucose and fructose) or with glucose only for each experimental condition. The theoretical monoculture (MC)-only yield is 0.6, so the presence of *Clj* in cocultures (CC) leads to that value being exceeded in serum bottles or to the monoculture yields being nearly doubled in bioreactors (BRs, 0.24-0.25 to 0.50-0.53 in high-SCD conditions).

| Letter | Yields considering glucose and fructose | Yields considering glucose |
| --- | --- | --- |
| Bottled MC – N<sub>2</sub> | 0.42 | 0.43 |
| Bottled CC – N<sub>2</sub> | 0.56 | 0.61 |
| Bottled CC – H<sub>2</sub> | 0.53 | 0.59 |
| Bottled CC – H<sub>2</sub>/CO<sub>2</sub> | 0.68 | 0.76 |
| Low-SCD MC BRs | 0.29 | 0.31 |
| High-SCD MC BRs | 0.24 | 0.25 |
| Low-SCD CC BRs | 0.46 | 0.48 |
| High-SCD CC BRs | 0.50 | 0.53 |

## Supplemental Figures

**Figure S1:**
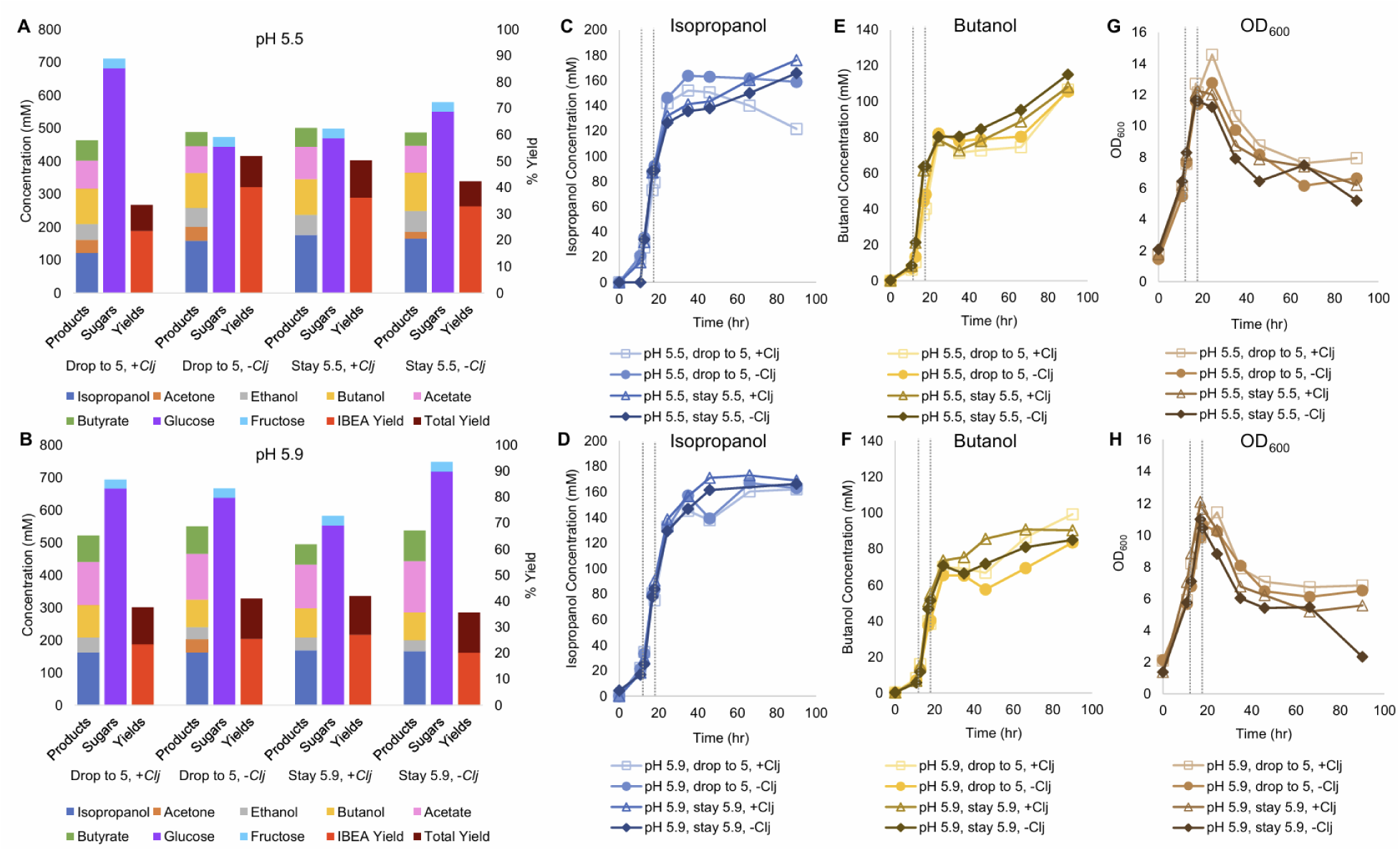
Metabolite production and yields, and sugar consumption to analyze pH setpoints and the impact of supplemental *Clj* cells for *Cac* and *Clj* cocultures in bioreactors. The vertical, short-dashed lines represent glucose additions to the bioreactors at 12 and 19 hours. (A-B) Summary of product concentrations, sugar consumption, and yields at a set pH of 5.5 (A) and at a set pH of 5.9 (B). IBEA yield refers only to IPA, BuOH, EtOH, and acetone. The total yield also includes butyrate and acetate. (C-D) The IPA formation kinetics of the pH 5.5 (C) and pH 5.9 (D) cocultures with (“drop to 5”) and without (“stay at”) a pH setpoint drop to 5.0 and additional *Clj* (*+Clj*; no additional *Clj* is *-Clj*). (E-F) The BuOH formation kinetics of the pH 5.5 (E) and pH 5.9 (F) cocultures. (G-H) The total biomass (OD_600_) of the pH 5.5 (G) and pH 5.9 (H) cocultures.

**Figure S2:**
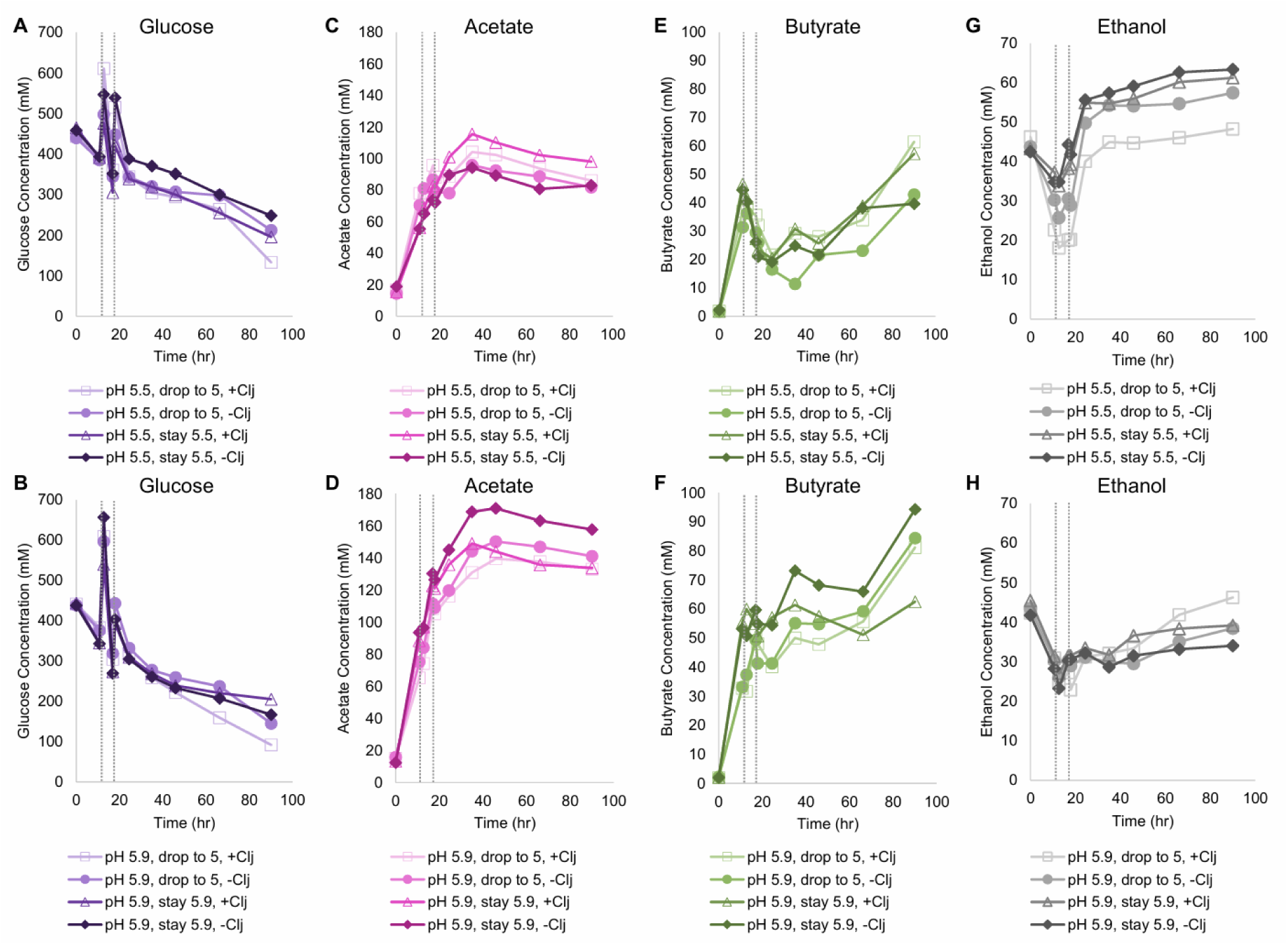
Kinetics of glucose, acetate, butyrate, and ethanol in fermentations to analyze pH setpoints and the impact of supplemental *Clj* cells for *Cac* and *Clj* cocultures in bioreactors. The vertical, short-dashed lines represent glucose additions to the bioreactors at 12 and 19 hours. (A-B) The glucose consumption kinetics of the pH 5.5 (A) and pH 5.9 (B) cocultures with (“drop to 5”) and without (“stay at”) a pH setpoint drop to 5.0 and additional *Clj* (*+Clj*; no additional *Clj* is *-Clj*). Glucose was fed twice at 12 and 19 hours. (C-D) Acetate kinetics of the pH 5.5 (C) and pH 5.9 (D) cocultures. (E-F) Butyrate kinetics at pH 5.5 (E) and pH 5.9 (F). (G-H) Ethanol kinetics at pH 5.5 (G) and pH 5.9 (H).

**Figure S3:**
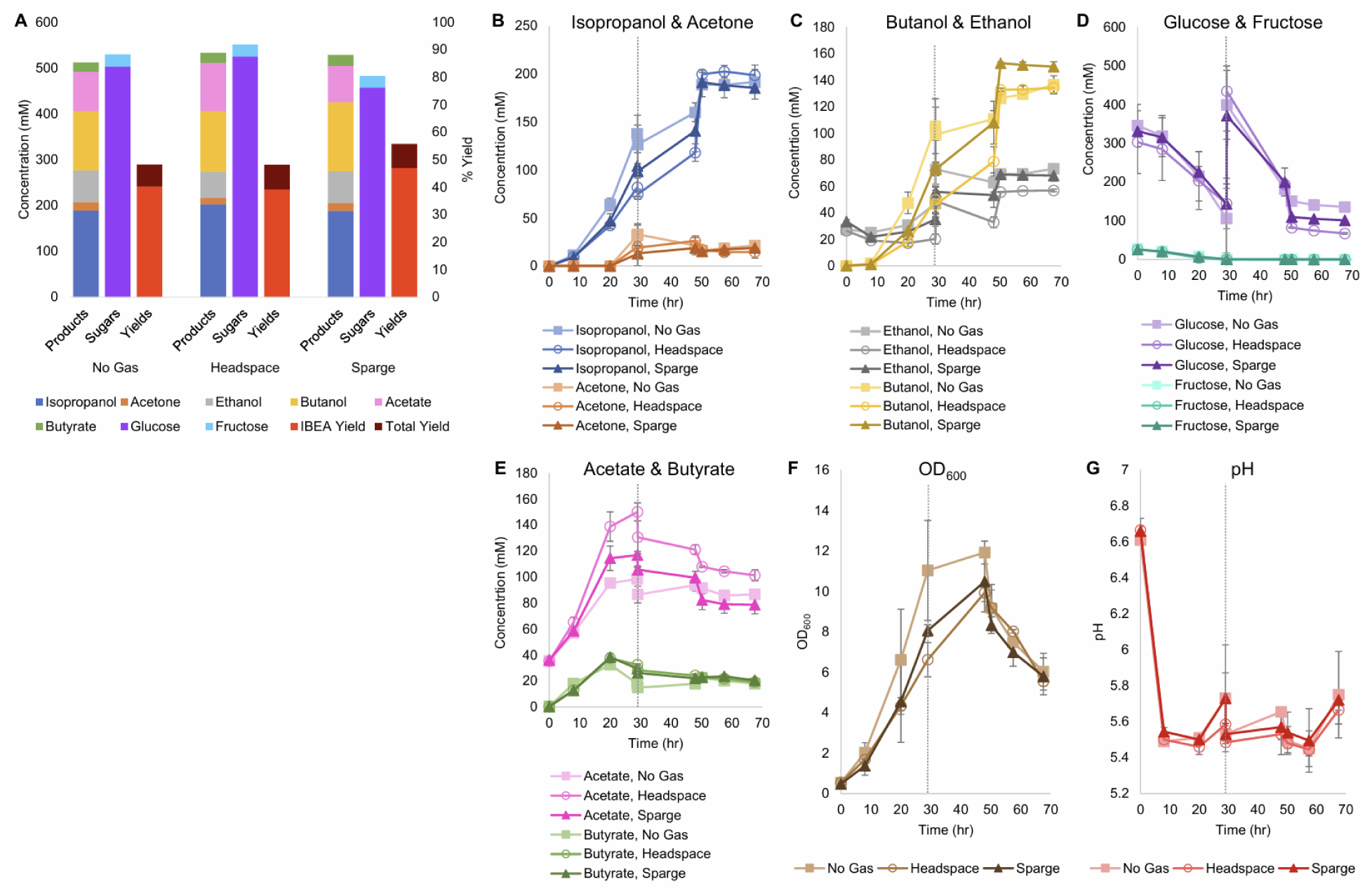
Metabolite production and yields, and sugar consumption, in bioreactor experiments to test gassing configurations for *Cac* and *Clj* cocultures. A blend 85% N_2_, 10% CO_2_, and 5% H_2_ was either sparged (bubbled) into the bioreactors, added to its headspace, or withheld (no gas) except during sampling. The vertical, short-dashed lines represent glucose additions to the bioreactors at 30 hours. (A) Summary of the metabolite concentrations and yields, and sugar consumption. IBEA yield refers to IPA, BuOH, EtOH, and acetone. The total yield also includes butyrate and acetate. (B) IPA and acetone concentration kinetics. (C) EtOH and BuOH formation kinetics. (D) Glucose and fructose consumption kinetics. (E) Acetate and butyrate concentration kinetics. (F) Biomass (OD_600_) kinetic profiles. (G) pH kinetic profiles.

**Figure S4:**
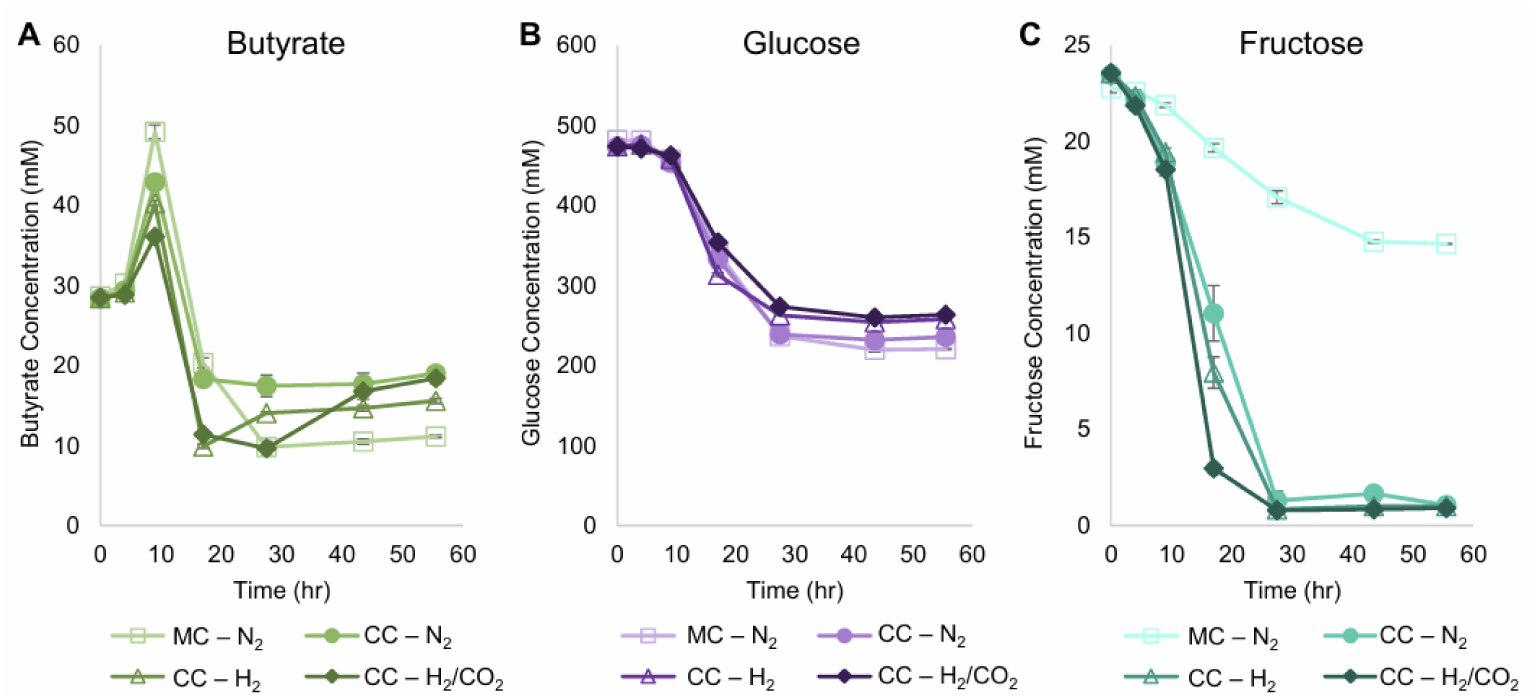
Butyrate and sugar kinetic profiles of monocultures (MC) and cocultures (CC) in sealed serum bottles (the experiments of Figure 2) with different headspace gas composition: N_2_, H_2_ or Mix [H_2_/CO_2_ (80/20)]. (A) Butyrate concentration kinetics. (B) Glucose consumption kinetics. (C) Fructose consumption kinetics.

**Figure S5:**
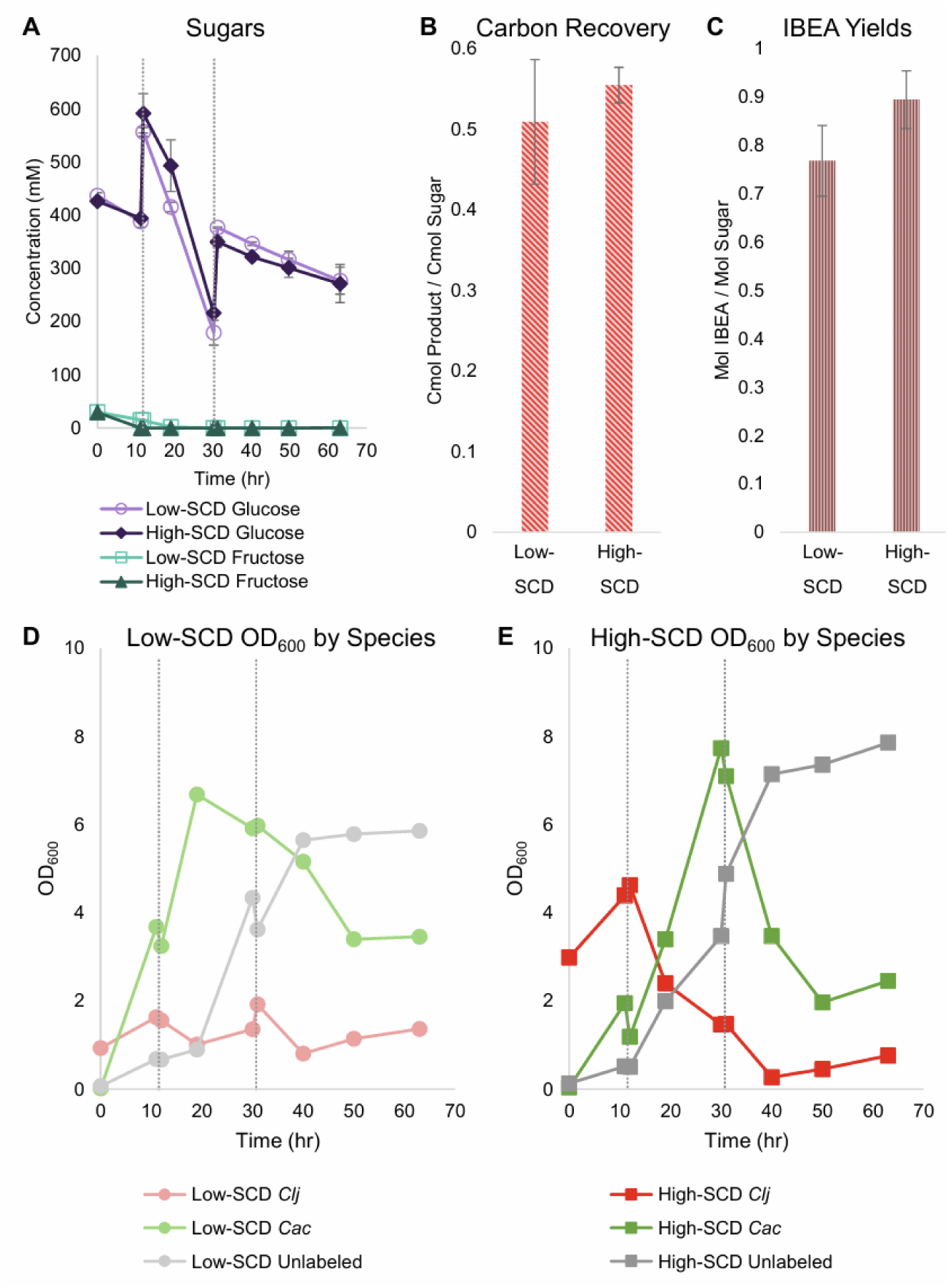
Sugar consumption, carbon recovery and IBEA yields, and biomass (OD_600_) of each species of the bioreactor cocultures of Figure 6 to assess the impact of low or high Starting Cell Densities (SCDs). The vertical, short-dashed lines represent glucose additions, at 10 and 30 hours, to the bioreactors. (A) Sugar consumption kinetics. (B) The carbon-moles (C-mols) of products produced per C-mols of sugars consumed. (C) Mol of alcohols (IBEA) produced per mols of sugar consumed. (D-E) The biomass (OD_600_) kinetics for each coculture species in the low-SCD (D) and high-SCD (E) bioreactors, as determined by the total OD_600_ and the percentage of each species determined by RNA-FISH labeling.

## Citations

Adams, M. R. (2017). “The birth of modern industrial microbiology: the acetone–butanol fermentation.” The international journal for the history of engineering & technology 87(1): 81–95.

Boy, C., J. Lesage, S. Alfenore, N. Gorret and S. E. Guillouet (2023). “Comparison of plasmid stabilization systems during heterologous isopropanol production in fed-batch bioreactor.” Journal of Biotechnology 366: 25–34.

Carlson, E. D. and E. T. Papoutsakis (2017). “Heterologous Expression of the Clostridium carboxidivorans CO Dehydrogenase Alone or Together with the Acetyl Coenzyme A Synthase Enables both Reduction of CO2 and Oxidation of CO by Clostridium acetobutylicum.” Appl Environ Microbiol 83(16): e00829.

Carrié, M., H. Velly, F. Ben-Chaabane and J.-C. Gabelle (2022). “Modeling fixed bed bioreactors for isopropanol and butanol production using Clostridium beijerinckii DSM 6423 immobilized on polyurethane foams.” Biochemical Engineering Journal 180: 108355.

Charubin, K., R. K. Bennett, A. G. Fast and E. T. Papoutsakis (2018). “Engineering Clostridium organisms as microbial cell-factories: challenges & opportunities.” Metab Eng 50: 173–191.

Charubin, K., J. D. Hill and E. T. Papoutsakis (2024). “DNA transfer between two different species mediated by heterologous cell fusion in Clostridium coculture.” mBio 15(2): e03133–03123.

Charubin, K., S. Modla, J. L. Caplan and E. T. Papoutsakis (2020). “Interspecies Microbial Fusion and Large-Scale Exchange of Cytoplasmic Proteins and RNA in a Syntrophic Clostridium Coculture.” mBio 11(5): e02030.

Charubin, K. and E. T. Papoutsakis (2019). “Direct cell-to-cell exchange of matter in a synthetic Clostridium syntrophy enables CO2 fixation, superior metabolite yields, and an expanded metabolic space.” Metab Eng 52: 9–19.

Cooksley, C. M., Y. Zhang, H. Wang, S. Redl, K. Winzer and N. P. Minton (2012). “Targeted mutagenesis of the Clostridium acetobutylicum acetone-butanol-ethanol fermentation pathway.” Metab Eng 14(6): 630–641.

Cui, Y., J. He, K.-L. Yang and K. Zhou (2020). “Aerobic acetone-butanol-isopropanol (ABI) fermentation through a co-culture of Clostridium beijerinckii G117 and recombinant Bacillus subtilis 1A1.” Metabolic engineering communications 11: e00137.

Dahle, M. L., E. T. Papoutsakis and M. R. Antoniewicz (2022). “13C-metabolic flux analysis of Clostridium ljungdahlii illuminates its core metabolism under mixotrophic culture conditions.” Metabolic Engineering 72: 161–170.

Dalal, J., M. Das, S. Joy, M. Yama and J. Rawat (2019). “Efficient isopropanol-butanol (IB) fermentation of rice straw hydrolysate by a newly isolated Clostridium beijerinckii strain C-01.” Biomass and Bioenergy 127: 105292.

Desai, R. P., L. K. Nielsen and E. T. Papoutsakis (1999). “Stoichiometric modeling of Clostridium acetobutylicum fermentations with non-linear constraints.” Journal of Biotechnology 71(1): 191–205.

Desai, R. P., L. K. Nielsen and E. T. Papoutsakis (1999). “Stoichiometric modeling of Clostridium acetobutylicum fermentations with non-linear constraints.” Journal of Biotechnology 71: 191–205.

Desai, R. P. and E. T. Papoutsakis (1999). “Antisense RNA Strategies for Metabolic Engineering of Clostridium acetobutylicum.” Applied and Environmental Microbiology 65(3): 936–945.

Dusseaux, S., C. Croux, P. Soucaille and I. Meynial-Salles (2013). “Metabolic engineering of Clostridium acetobutylicum ATCC 824 for the high-yield production of a biofuel composed of an isopropanol/butanol/ethanol mixture.” Metab Eng 18: 1–8.

Foulquier, C., A. Rivière, M. Heulot, S. Dos Reis, C. Perdu, L. Girbal, M. Pinault, S. Dusséaux, M. Yoo, P. Soucaille and I. Meynial-Salles (2022). “Molecular characterization of the missing electron pathways for butanol synthesis in Clostridium acetobutylicum.” Nature Communications 13(1): 4691.

Gabriel, C. and F. Crawford (1930). “Development of the Butyl-Acetonic Fermentation Industry.” Industrial and Engineering Chemistry 22(11): 1163–1165.

Gibbs, D. (1983). “The rise and fall (…and rise?) of acetone/butanol fermentations.” Trends in Biotechnology 1: 12–15.

Girbal, L., C. Croux, I. Vasconcelos and P. Soucaille (1995). “Regulation of metabolic shifts in Clostridium acetobutylicum ATCC 824.” FEMS Microbiology Reviews 17(3): 287–297.

Grimmler, C., H. Janssen, D. Krauβe, R.-J. Fischer, H. Bahl, P. Dürre, W. Liebl and A. Ehrenreich (2011). “Genome-Wide Gene Expression Analysis of the Switch between Acidogenesis and Solventogenesis in Continuous Cultures of Clostridium acetobutylicum.” Microbial Physiology 20(1): 1–15.

Grupe, H. and G. Gottschalk (1992). “Physiological Events in Clostridium acetobutylicum during the Shift from Acidogenesis to Solventogenesis in Continuous Culture and Presentation of a Model for Shift Induction.” Applied and Environmental Microbiology 58(12): 3896–3902.

Haus, S., S. Jabbari, T. Millat, H. Janssen, R.-J. Fischer, H. Bahl, J. R. King and O. Wolkenhauer (2011). “A systems biology approach to investigate the effect of pH-induced gene regulation on solvent production by Clostridium acetobutylicum in continuous culture.” BMC Systems Biology 5(1): 10.

Hill, J. D. and E. T. Papoutsakis (2024). “Species-specific ribosomal RNA-FISH identifies interspecies cellular-material exchange, active-cell population dynamics and cellular localization of translation machinery in clostridial cultures and co-cultures.” mSystems 9(10): e00572–00524.

Hocq, R. and M. Sauer (2022). “An artificial coculture fermentation system for industrial propanol production.” FEMS Microbes 3: xtac013.

Inokuma, K., J. C. Liao, M. Okamoto and T. Hanai (2010). “Improvement of isopropanol production by metabolically engineered Escherichia coli using gas stripping.” J Biosci Bioeng 110(6): 696–701.

Jiang, Y., R. Wu, W. Zhang, F. Xin and M. Jiang (2023). “Construction of stable microbial consortia for effective biochemical synthesis.” Trends in biotechnology (Regular ed.) 41(11): 1430–1441.

Jojima, T., M. Inui and H. Yukawa (2008). “Production of isopropanol by metabolically engineered Escherichia coli.” Applied Microbiology and Biotechnology 77(6): 1219–1224.

Jones, S. W., A. G. Fast, E. D. Carlson, C. A. Wiedel, J. Au, M. R. Antoniewicz, E. T. Papoutsakis and B. P. Tracy (2016). “CO2 fixation by anaerobic non-photosynthetic mixotrophy for improved carbon conversion.” Nature Communications 7(1): 12800.

Katsyv, A. and V. Müller (2020). “Overcoming energetic barriers in acetogenic C1 conversion.” Frontiers in Bioengineering and Biotechnology 8: 621166.

Lee, J., Y.-S. Jang, S. J. Choi, J. A. Im, H. Song, J. H. Cho, D. Y. Seung, E. T. Papoutsakis, G. N. Bennett and S. Y. Lee (2012). “Metabolic Engineering of Clostridium acetobutylicum ATCC 824 for Isopropanol-Butanol-Ethanol Fermentation.” Applied and Environmental Microbiology 78(5): 1416–1423.

Lee, J. Y., Y.-S. Jang, J. Lee, E. T. Papoutsakis and S. Y. Lee (2009). “Metabolic engineering of Clostridium acetobutylicum M5 for highly selective butanol production.” Biotechnology journal 4(10): 1432–1440.

Lutke-Eversloh, T. and H. Bahl (2011). “Metabolic engineering of Clostridium acetobutylicum: recent advances to improve butanol production.” Curr Opin Biotechnol 22(5): 634–647.

Mermelstein, L. D. and E. T. Papoutsakis (1993). “In vivo methylation in Escherichia coli by the Bacillus subtilis phage phi 3T I methyltransferase to protect plasmids from restriction upon transformation of Clostridium acetobutylicum ATCC 824.” Applied and Environmental Microbiology 59(4): 1077–1081.

Moon, Y. H., K. J. Han, D. Kim and D. F. Day (2015). “Enhanced production of butanol and isopropanol from sugarcane molasses using Clostridium beijerinckii optinoii.” Biotechnology and Bioprocess Engineering 20(5): 871–877.

Otten, J. K., Y. Zou and E. T. Papoutsakis (2022). “The potential of caproate (hexanoate) production using Clostridium kluyveri syntrophic cocultures with Clostridium acetobutylicum or Clostridium saccharolyticum.” Front Bioeng Biotechnol 10: 965614.

Papoutsakis, E. T. (1984). “Equations and calculations for fermentations of butyric acid bacteria.” Biotechnology and bioengineering 26(2): 174–187.

Procentese, A., F. Raganati, L. Navarini, G. Olivieri, M. E. Russo and A. Marzoccchella (2018). “Coffee Silverskin as a Renewable Resource to Produce Butanol and Isopropanol.” Chemical Engineering Translations 64: 139–144.

Richter, H., B. Molitor, M. Diender, D. Z. Sousa and L. T. Angenent (2016). “A Narrow pH Range Supports Butanol, Hexanol, and Octanol Production from Syngas in a Continuous Co-culture of Clostridium ljungdahlii and Clostridium kluyveri with In-Line Product Extraction.” Front Microbiol 7: 1773.

Rochón, E., F. Cebreiros, M. D. Ferrari and C. Lareo (2019). “Isopropanol-butanol production from sugarcane and sugarcane-sweet sorghum juices by Clostridium beijerinckii DSM 6423.” Biomass and Bioenergy 128: 105331.

Seo, H., S. H. Capece, J. D. Hill, J. K. Otten and E. T. Papoutsakis (2024). “Butyrate as a growth factor of Clostridium acetobutylicum.” Metabolic Engineering 86: 194–207.

Sillers, R., M. A. Al-Hinai and E. T. Papoutsakis (2009). “Aldehyde-alcohol dehydrogenase and/or thiolase overexpression coupled with CoA transferase downregulation lead to higher alcohol titers and selectivity in Clostridium acetobutylicum fermentations.” Biotechnol Bioeng 102(1): 38–49.

Streett, H. E., K. M. Kalis and E. T. Papoutsakis (2019). “A Strongly Fluorescing Anaerobic Reporter and Protein-Tagging System for Clostridium Organisms Based on the Fluorescence-Activating and Absorption-Shifting Tag Protein (FAST).” Applied and Environmental Microbiology 85(14): e00622–00619.

Willis, N. B., J. K. Otten, H. Seo, P. C. Munasinghe, J. D. Hill and E. T. Papoutsakis (2025). “Enabling supratheoretical isopropanol yields from carbon-negative glucose fermentations with Clostridium acetobutylicum-Clostridium ljungdahlii cocultures.” bioRxiv: 10.1101/2025.1107.1114.664808.

Willis, N. B. and E. T. Papoutsakis (2025). “Separate, separated, and together: the transcriptional program of the Clostridium acetobutylicum-Clostridium ljungdahlii syntrophy leading to interspecies cell fusion.” Msystems 10(5): e00030–00025.

Zhang, F., K. Zhang, Z. Zhang, H.-Q. Chen, X.-W. Chen, X.-Y. Xian and Y.-R. Wu (2023). “Efficient isopropanol-butanol-ethanol (IBE) fermentation by a gene-modified solventogenic Clostridium species under the co-utilization of Fe(III) and butyrate.” Bioresource Technology 373: 128751.

