## Appendix 1 for "Cross-talk between engineered *Clostridium acetobutylicum* and *Clostridium ljungdahlii* in syntrophic cocultures enhances isopropanol and butanol production"

### Appendix 1: Stoichiometric Coefficient Calculations and Details

#### Chemical Species in CC Model

**Extracellular metabolites: (accumulation determined by HPLC)**

1. glucose
2. fructose
3. acetate
4. ethanol
5. isopropanol
6. butyrate
7. acetone
8. butanol

***C. acetobutylicum* intracellular metabolites: (assumed to not accumulate)**

9. pyruvate-(Cac)
10. acetyl-CoA-(Cac)
11. acetoacetyl-CoA-(Cac)
12. butyryl-CoA-(Cac)
13. NADH-(Cac)
14. $Fd_{red}$-(Cac)

***C. ljungdahlii* intracellular metabolites: (assumed to not accumulate)**

15. pyruvate-(Clj)
16. acetyl-CoA-(Clj)
17. EC-(Clj)

#### Reactions in the CC Model

**Reaction 1: rGLY1.Cac**

glucose + 2 $NAD^+$-(Cac) → 2 pyruvate-(Cac) + 2 NADH-(Cac)

**Reaction 2: rGLY2.Cac**

pyruvate-(Cac) → acetyl-CoA-(Cac) + $Fd_{red}$-(Cac) + $CO_2$

**Reaction 3: rFDNH**

$Fd_{red}$-(Cac) + $NAD^+$-(Cac) → $Fd_{ox}$-(Cac) + NADH-(Cac)

**Reaction 4: rHYD.Cac**

$Fd_{red}$-(Cac) + 2 $H^+$ → $Fd_{ox}$-(Cac) + $H_2$

**Reaction 5: rPTAAK.Cac**

acetyl-CoA-(Cac) → acetate

**Reaction 6: rETOH**

acetyl-CoA-(Cac) + 2 NADH-(Cac) → ethanol + 2 $NAD^+$-(Cac)

**Reaction 7: rTHL**

2 acetyl-CoA-(Cac) → acetoacetyl-CoA-(Cac)

**Reaction 8: rBYCA**

acetoacetyl-CoA-(Cac) + $Fd_{ox}$-(Cac) + 3 NADH-(Cac) → butyryl-CoA-(Cac) + $Fd_{red}$-(Cac) + 3 $NAD^+$-(Cac)

**Reaction 9: rPTBBK**

butyryl-CoA-(Cac) → butyrate

**Reaction 10: rBUOH**

butyryl-CoA-(Cac) + 2 NADH-(Cac) → butanol + 2 $NAD^+$-(Cac)

**Reaction 11: rBYUP**

butyrate + acetoacetyl-CoA-(Cac) → acetone + butyryl-CoA-(Cac) + $CO_2$

**Reaction 12: rACUP**

acetate + acetoacetyl-CoA-(Cac) → acetone + acetyl-CoA-(Cac) + $CO_2$

**Reaction 13: rGLY1.Clj**

fructose + 2 $EC_{ox}$-(Clj) → 2 pyruvate-(Clj) + 2 $EC_{red}$-(Clj)

**Reaction 14: rGLY2.Clj**

pyruvate-(Clj) + $EC_{ox}$-(Clj) → acetyl-CoA-(Clj) + $EC_{red}$-(Clj) + $CO_2$

**Reaction 15: rPTAAK.Clj**

acetyl-CoA-(Clj) → acetate

**Reaction 16: rHYD.Clj**

$H_2$ + $EC_{ox}$-(Clj) → 2 $H^+$ + $EC_{red}$-(Clj)

**Reaction 17: rWLP**

2 $CO_2$ + 4 $EC_{red}$-(Clj) → acetyl-CoA-(Clj) + 4 $EC_{ox}$-(Clj)

**Reaction 18: rSADH**

acetone + $EC_{red}$-(Clj) → isopropanol + $EC_{ox}$-(Clj)

**Details**
The specific intracellular Electron Carrier ($EC_{ox}/EC_{red}$) in *Clj* is unimportant because the 'rnf/ATPase' complex allows interconversion between NADH and $Fd_{red}$ at the expense of ATP, which is not accounted for. This is in contrast to *Cac*'s electron carriers which are specified as being $NAD^+$/NADH or $Fd_{ox}/Fd_{red}$.

$CO_2$ and $H_2$ are included in the above equations for convenience, but their accumulation cannot be used to constrain the model's solution because their accumulation was not measured.

In theory, this is an under-defined matrix, because there are 18 unique reactions, and only 17 metabolites. Therefor it could generally find a 'perfect' solution, where the sum of the square residuals (SSR) is 0. However, that may involve running certain reactions in reverse, which may or may not be physiologically possible.  In practice, I use the non-negative least squares (nnls) solver in SciPy, such that no element of the x-matrix can be less than 0. This means that the solution space may not always contain a particular solution in which the SSR is 0. When this occurs, the solver relies on a minimum finding algorithm to find the particular solution with the smallest possible SSR.
